# Bloom-forming bacteria heavily invest in anti-phage defense

**DOI:** 10.64898/2026.09.26.754401

**Authors:** Roo Weed, Molly A Moynihan, Eleanor Greene, Olivia L Mathieson, Catherine A Crowley, Emily N Junkins, Hannah Vanderscheuren, Abdeali M Jivaji, Manuel Kleiner, S Emil Ruff

**Author notes:** Present addresses: EG: University of Amsterdam, Netherlands, CAC: University of Connecticut, Groton, CT, USA, ENJ: Nantucket Conservation Foundation, Nantucket, MA, 02554, USA.

## Abstract

Bacterial blooms are characterized by unusually high cell densities and exceptionally low diversity and can profoundly alter ecosystem function and services. Bacteriophages have long been considered an important cause of mortality in blooms, acting as a mechanism for control. Here, we characterize the viral ecology of a long-lasting estuarine bloom of green sulfur bacteria (Chlorobiota). We combined direct cell and viral counts with metagenomic and metaproteomic data to characterize host and phage activity at different time points. The abundance of virus-like particles (VLPs) decreased at high cell densities, suggesting reduced lytic infection rates. The dominant organism, GSB-TRL01 (genus *Prosthecochloris*), apparently contained a large conjugative plasmid encoding five different anti-phage defense systems. The organism’s genome encoded 13 additional defense systems. Compared to the average of five defense systems per microbial genome, this enrichment suggests robust anti-phage defense capabilities. Proteins from ten different defense systems on GSB-TRL01’s genome and four systems from the conjugative plasmid were expressed in the proteome. This suggests that GSB-TRL01 invests heavily in anti-phage defense, leading to reduced lysis at high cell densities and allowing blooms to persist for weeks to months. To determine whether this ability is widespread among bloom forming organisms, we compared genomes of putative bloomers to those of non-blooming organisms. We found that bloomer genomes were significantly enriched with anti-phage defense systems. This challenges traditional paradigms of phage ecology in bloom-forming systems and suggests that microbes adapted to high-density growth may have evolved mechanisms to reduce their susceptibility to phage attack.

## Introduction

Bacterial blooms are dense assemblages of one or a few bacterial populations that grow to exceptionally high cell densities over short time scales, profoundly altering ecosystem functions and services. Blooms are widespread and occur in diverse ecosystems such as stratified meromictic lakes [1–3], the open ocean [4, 5], and coastal systems [6, 7]. Striking blooms of anoxygenic phototrophs occur yearly in the Trunk River Lagoon (TRL) estuary in Falmouth, MA (N 41.535236, W −70.641298) [7, 8]. TRL is a brackish system connected to Vineyard Sound and Oyster Pond (figure 1a), and is therefore influenced by tides, storms, and runoff.

**Figure 1.**
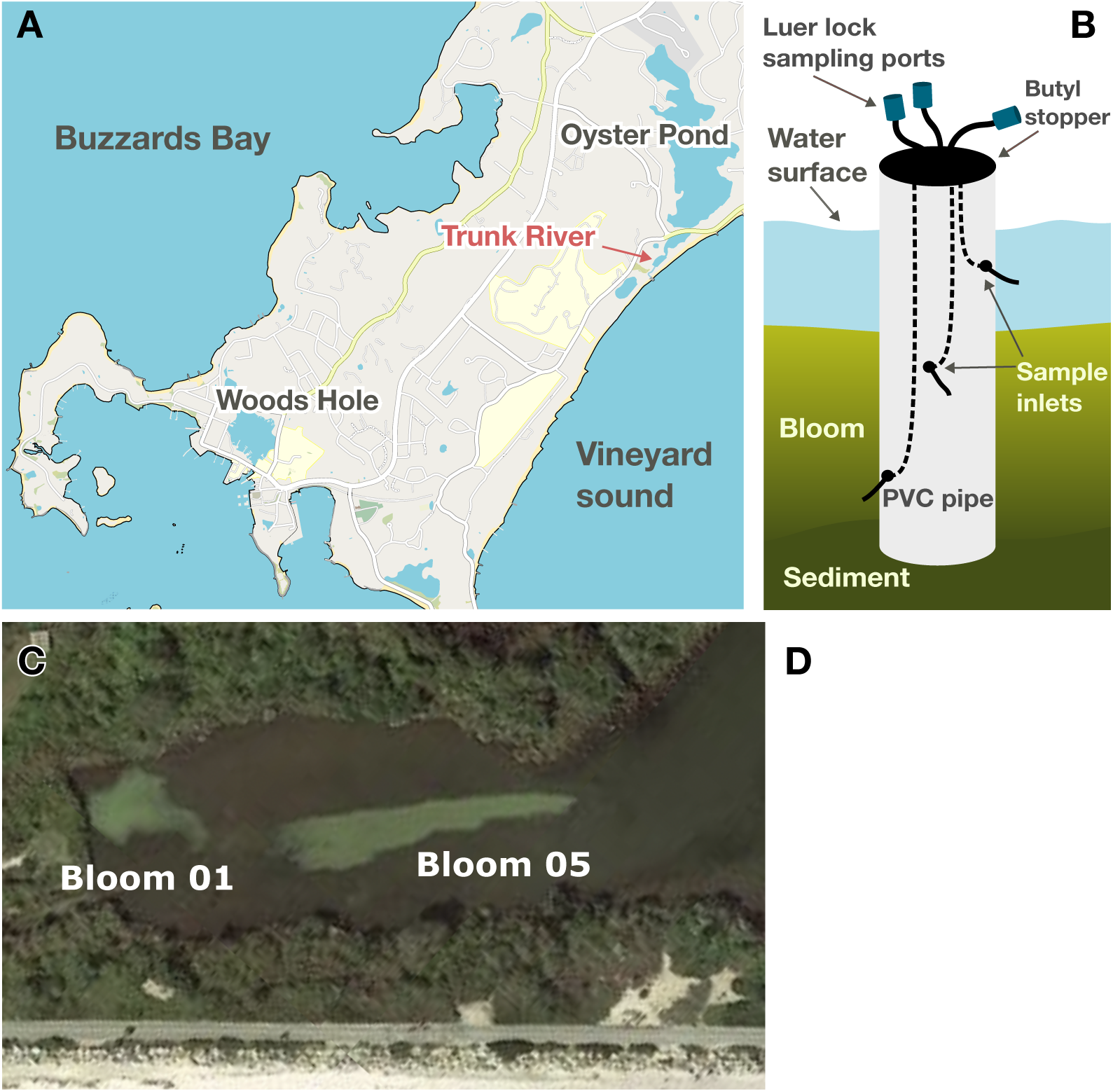
Trunk River Lagoon location and bloom sampling. a. Map of the southern tip of Cape Cod, MA, USA, b. diagram and photo of the sampling pole, c. Satellite photograph of Trunk River from Google Earth, taken on October 6, 2018, d. Photograph showing a bloom on August 26th, 2021.

Stratification between the upper oxic water column and the lower euxinic layer provides a niche for sulfur-oxidizing anoxygenic phototrophic green sulfur bacteria (genus *Prosthecochloris*) which frequently bloom during the summer months [8]. Blooms form between 30 and 60 cm below the water surface and extend to the sediment interface at (70-100 cm depth), with the highest cell densities towards the top of the bloom (figure 1b). TRL blooms are characterized by extremely high cell densities (up to ∼10^9^ cells mL^-1^), persistence (lasting weeks to months), and near clonality [8], and represent an excellent model system in which to characterize phage infection and resistance in a bloom context.

Bacteriophages are suggested to be important agents of mortality in bacterial blooms and have been proposed as possible tools for bloom control [9, 10]. Yet phage ecology in bacterial blooms generally remains poorly understood.

A prevailing view of viral ecology in bloom environments has been that blooming organisms “boom” and are often subsequently “busted” by phages, thus implicating phages in bloom decline [11–15]. However, ratios of virus-like particles (VLP) to cells have been observed to decline at high cell densities in anoxygenic phototrophic bloom systems [1, 3, 16] pointing towards a need to reconsider the “boom and bust” paradigm.

One of the dominant theoretical frameworks in phage ecology is the Kill-the-Winner (KtW) hypothesis, which is often understood as predicting “boom-and-bust” dynamics in blooms [14, 17]. In fact, because predation pressure is density dependent [18], KtW predicts that faster growing “winners” (“competition specialists”), will be targeted by phage attack, thereby allowing slower growing “defense specialists” to compete, and diversifying the overall community [19–22]. KtW therefore predicts that under nutrient replete conditions, such as those in TRL, communities may be dominated by defense specialists, which succeed despite the potential costs of resistance in optimal conditions [23]. The prevalence of anti-phage defense related genes in the genomes of some bloom forming Cyanobacteria [24–27] suggests that the accumulation of anti-phage defense systems may occur in organisms which can maintain high-density populations. The accumulation of anti-phage defense genes would be supported by the high prevalence of horizontal gene transfer in high-density populations (the “King of the Mountain” hypothesis) [28].

We hypothesized that the anoxygenic phototrophic bacteria that bloom in TRL, and indeed bacterial bloom formers in general, are defense specialists, accumulating high numbers of anti-phage defense systems that allow them to defend against diverse phage attacks. Using datasets from 2021, 2024 and 2025, we characterized the viral ecology in TRL blooms with the following objectives: 1) determine the abundance dynamics of viral particles; 2) identify phages that likely infect the dominant bloom-forming organism; 3) understand the anti-phage defense systems employed by GSB-TRL01 and an associated conjugative plasmid to counter these attacks. We generalized our findings by comparing the abundance of anti-phage defense systems in the genomes of putative bloom forming bacteria with non-bloomers to determine whether investment in anti-phage defense systems is a common ecological strategy among high-density adapted microbial populations.

## Materials and methods

### Study Site and Sample collection

Our data include samples from three different years in TRL. We collected samples using a custom PVC sampling pole (figure 1b, [7, 8]), which allowed sample collection from consistent depths in the water column over time. In 2021, samples were collected as in [8] from blooms in two distinct areas of the estuary referred to as Bloom 01 (B01) and Bloom 05 (B05) (figure 1c). Samples were collected over seven consecutive days from 26 August 2021 to 1 September 2021 and then again on 10 September 2021, and finally on 20 and 21 September 2021. The bloom in the B05 area lasted 56 days, whereas the bloom in the B01 area lasted only 4 days. We sampled for direct cell counts, metagenomics, and metaproteomics (Table 1, supplementary table 1). Short-read metagenomics samples were collected using sequential filtration (pore sizes: 2 µm, 0.2 µm, 0.1 µm, and 0.025 µm; MCE Membrane Filters, Millipore Sigma), generating size fractions of 0.2-2.0 µm, 0.1-0.2 µm, and 0.025-0.1 µm. We collected the sample for long-read Nanopore sequencing on a 0.22 μm polyethersulfone filter (Sterivex, SVGP01050). For metaproteomics, we collected a cellular fraction by filtering the bloom onto a 0.22 μm Sterivex filter and an extracellular fraction by centrifuging samples at 20,000 × rcf for 15 minutes at 4°C and retaining the supernatant.

**Table 1.** An outline of the samples collected for direct counts, metagenomics, and metaproteomics for each year. Supplementary table 1 contains a full description of all samples collected.

|  | 2021 | 2024 | 2025 |
| --- | --- | --- | --- |
| **DAPI cell counts** | 11 | 50 | - |
| **CARD-FISH cell counts** | 4 | - | - |
| **Sybr green viral counts** | - | 50 | - |
| **> 0.2 µm size fraction metagenomics** | 1<br>(Nanopore sequencing) | 2 | 1 |
| **0.2-2.0 µm size fraction metagenomics** | 3 | - | - |
| **0.1-0.2 µm size fraction metagenomics** | 5 | - | - |
| **0.025-0.1 µm size fraction metagenomics** | 6 | - | - |
| **Intracellular metaproteomic fraction** | 15 | - | - |
| **Extracellular metaproteomic fraction** | 12 | - | - |

In 2024, samples were collected approximately weekly from 14 June 2024 to 20 August 2024 at five depths in the same bloom in the B05 area (figure 1c). The bloom lasted a total of 55 days. From this sampling period, we include direct cell and VLP counts, and metagenomic samples (Table 1, supplementary table 1). On 9 September 2025 we collected one sample for metagenomics from a bloom in the B05 area. All omics samples were stored at −80 °C until processed.

### Direct cell and virus-like particle (VLP) counts

We performed direct cell counts as described in [8]. A detailed description of the protocol for cell and VLP enumeration is available in the supplementary materials. Briefly, we filtered fixed cells onto 0.2 µm filters, stained them with DAPI (4’,6-diamidino-2-phenylindole), and enumerated them with epifluorescence microscopy. We performed direct VLP counts as described in [29]. Briefly, we pre-filtered samples with a 0.2 µm filter to remove cells, fixed the filtrate, filtered fixed particles onto a 0.02 µm filter, stained with Sybr Green I, and enumerated VLPs with epifluorescence microscopy.

### Catalyzed reporter deposition fluorescence in situ hybridization (CARD-FISH)

CARD-FISH was performed as in [8]. A detailed description of the protocol is available in the supplementary materials. Briefly, we used two probes: GSB-532, targeting green sulfur bacteria [30] and NON338, a nonsense probe [31]. We fixed the cells in 2% formaldehyde in 1× PBS, filtered samples onto 0.2 μm polycarbonate filters using 0.45 μm nitrocellulose support filters, permeabilized cells with lysozyme, incubated the cells with a hybridization buffer containing either one of the oligonucleotide probes (50 ng μl^-1^ f.c. for GSB-532; 0.16 ng μl^-1^ f.c. for NON338), and performed signal amplification with the fluorescent dye Alexa Fluor 488 (Invitrogen, B40957).

### DNA extraction

DNA extraction was performed as in [8]. A detailed protocol for DNA extraction is available in the supplementary materials. Briefly, we performed manual DNA extractions with a CTAB-based extraction buffer using mechanical (freeze-thaw cycles), chemical (Sodium dodecyl sulfate) and enzymatic (lysozyme, Proteinase K) lysis steps. We isolated DNA using chloroform-isoamyl alcohol and precipitated using 100% isopropanol. To quantify DNA, we used a high-sensitivity Qubit DNA assay and a NanoDrop 2000 spectrophotometer.

### Oxford Nanopore Sequencing

We sequenced extracted DNA (>0.2 µm size fraction) with Oxford Nanopore Technologies (ONT) ligation sequencing kit (SQK-LSK110) following the manufacturer’s instructions. A detailed protocol for DNA extraction is available in the supplementary materials. We loaded the resulting library (8.9 ng/µL) onto the flow cell (FLOW-MIN106) and sequenced with a MinION sequencer (Mk1B; ONT) following the SQKLSK110 manufacturer’s protocol.

### Nanopore Sequence Data Processing

We processed Nanopore sequence data as described [32], with recommendations for assembling bacterial genomes from Oxford Nanopore sequencing with short-read Illumina polishing. A detailed protocol is available in the supplementary materials. Briefly, we converted fast5 files to pod5 format (pod5 v0.3.6 Python module, ONT), basecalled files with Dorado (v0.5.3, ONT), quality filtered (Fitlong v0.2.1, [33]), and assembled reads (Flye v2.9.3, [34]). We then sequence corrected (Medaka v1.6.0, ONT) and polished long-reads with the Illumina short reads (Polypolish v0.6.0), [35]). The resulting assembly produced 9 contigs, one of which was the 2.2 Mb genome of GSB-TRL01 (determined to be circular by Flye), and another 92 kb long circular contig which belongs to a conjugative plasmid putatively associated with GSB-TRL01. We annotated the 2.2 Mb contig by manually combining annotations in regions of interest from Prokka v1.14.5 [36], Cenote-Taker 3 v3.0.0 [37], and DefenseFinder v1.2.2 [38]. The phage satellite was classified using SatelliteFinder [39]. Sequences were visualized using Geneious Prime 2025.1.3 (https://www.geneious.com) and Gene Graphics [40].

### Illumina sequencing

Shotgun metagenomics was performed by Psomagen Inc, as described [8]. A detailed protocol is available in the supplementary materials. Briefly, the Illumina DNA Prep Kit was used to prepare libraries and tagmented samples were amplified by PCR with an Illumina Nextera DNA Unique Dual Indexes kit. Libraries were sequenced on a NovaSeq6000 S4 (v1.5) platform with PE150 chemistry (paired-end sequencing, 150 bp).

### Short read metagenomic data processing and metagenome assembled genome (MAG) generation

We performed read quality control and generated MAGs as described [8]. Briefly, we removed adapters using trimmomatic and removed low quality reads using iu-filter-quality-minoche from illumina-utils (v2.12) [41]. We used MEGAHIT (v1.2.9) [42] to assemble quality controlled reads, then performed binning with four different binning algorithms and combined the output into a set of optimized, non-redundant bins. Only metagenomic samples from 2021 were included in the assemblies.

To create the initial bin database, we selected optimized and non-redundant bins from B05 samples only and included the 2.2 Mb contig from the Nanopore assembly. To assess bin quality, we used CheckM2 v1.0.1 [43] and retained only medium quality and up bins (≥50% completion, ≤10% contamination, standards from [44]). We further dereplicated bins using dRep v3.6.2 [45] with default settings (ANI >95%, <75% completeness and >25% contamination).

This resulted in a final database containing 65 dereplicated MAGs. We used gtdbtk de_novo_wf v2.4.1 [46] to determine taxonomy, with p Chloroflexota as the outgroup for bacterial taxonomic ID and p Undinarchaeota as the outgroup for Archaeal taxonomic ID.

### Identification of viral sequences and viral metagenome assembled genome (vMAG) generation

The full viral sequence identification and vMAG generation protocol is described in the supplementary methods. Briefly, we combined quality controlled reads by size fraction (0.025-0.1 µm and 0.1-0.22 µm) across all time points and assembled them using MEGAHIT v1.2.9 [42]. To identify contigs as viral, we used geNomad v1.8.1 [47] and VirSorter2 v2.2.4 [48], then used CheckV v1.0.1 [49] to assess quality and manually filtered the contigs according to standards modified from [50]. We dereplicated quality controlled contigs and binned them using vRhyme v1.1.0 [51]. Only vMAGs longer than 10 kbp and classified as “High quality” or “Complete” by CheckV were used in downstream analyses. We assigned taxonomy with geNomad [47] and ViralRecall v3.0 (https://github.com/abdealijivaji/ViralRecall_3.0) [52]. We used Cenote-Taker 3 v3.0.0 [37] for functional annotations and CoverM v0.6.1 [53] for relative abundance calculations for both MAGs and vMAGs.

### vMAG host identification

We used iPHoP v1.3.3 [54] with the Aug_2023_pub_rw database modified to include the prokaryotic bin database described above (yet un-dereplicated) to identify putative hosts of vMAGs. When a vMAG was matched to multiple hosts, we selected the CRISPR or Blast hit with the highest confidence. If only iPHoP-RF was available, then we selected the score with the highest iPHoP-RF confidence.

### Metaproteomics

Metaproteomics analysis was conducted as described [8]. A detailed description for each sample type is available in the supplementary methods. Briefly, for the cellular fraction we prepared lysates from cell material collected on Sterivex filters by suspending and heating the filters in a lysis buffer (4% w/v SDS, 100 mM Tris-HCl pH 8.0) as previously described [55]. For the extracellular fraction, we concentrated the supernatants of pelleted samples before suspending them in lysis buffer and heating. We precipitated proteins overnight with 10% trichloroacetic acid at −80 °C, washed the pellets with ice cold acetone and allowed the pellets to air dry. We suspended and heated protein precipitates in 60-120 µl of 4% (w/v) SDS, 100 mM Tris-HCl pH 7.6, 0.1 M dithiothreitol (DTT), before proceeding with a modified filter-aided sample preparation method to prepare for LC-MS/MS analysis [56].

We loaded 2000 ng of peptide onto an UltiMate 3000 RSLCnano UHPLC system (Thermo Fisher Scientific) for separation. Samples were first loaded onto a 5 mm, 500 μm C18 Acclaim PepMap100 pre-column (Thermo Fisher Scientific) for desalting, prior to separation on a 75 μm x 75 cm EASY-spray column packed with PepMap RSLC C18, 2 μm material (Thermo Fisher Scientific) at 60°C. We separated peptides using a previously published 140 minute reverse-phase gradient [57]. The eluting peptides were ionized via electrospray ionization (ESI) with an Easy-Spray source and measured using an Orbitrap Exploris 480 Mass Spectrometer (Thermo Fisher Scientific) by a data dependent acquisition [57].

We created a protein sequence database using predicted protein sequences from the MAGs, vMAGs, and unbinned sequences generated by the Illumina short-read assembly from 2021 following previously described procedures [58]. The Nanopore sequence data was later matched to this database using mmseqs easy-search [59]. We required sequences to have >95% amino acid percent identity to be considered a match. We searched the raw spectral data against our protein sequence database with Proteome Discoverer version 2.3 (Thermo Fisher Scientific) as previously described [60]. Normalized Spectral Abundance Factor percent (NSAF%) values for GSB-TRL01 and the associated plasmid have been normalized to represent the within-organism relative abundance only (orgNSAF%) [61–63].

### Broader analysis of bloom forming prokaryotes

To create a list of prokaryotic organisms thought to form blooms, we conducted an extensive literature review (supplementary table 8). We estimated the species-specific cell count using available data (ideally organism specific direct counts, e.g. via CARD-FISH, but when these were not available, we combined direct counts with sequence data) and considered an organism to be a bloom former if the count exceeded 80,000 cells mL^-1^ [64]. To capture dominant organisms without corresponding direct counts, we additionally included organisms with ≥ 50% relative abundance in metagenomic samples. To do this, we filtered the Sandpiper database [65] and only retained organisms with a species level taxonomic ID and relative abundance ≥ 50% in one or more samples with prokaryotic fraction greater than 50% in an aquatic environment (see supplementary methods for the full list of categories).

Organisms recovered from this filtering were manually checked to confirm that they had relative abundance ≥ 50% in a sample that was genuinely an environmental metagenome, not an experimental system or enrichment. Metagenomes from sediment were excluded due to the difficulty of determining sedimentary cell densities. Metagenomes from hot springs were also excluded because prokaryotes from hot springs have previously been shown to encode high numbers of anti-phage defense systems [66], likely due to eco-evolutionary forces outside the scope of this project.

The final database contained 89 putative bloom forming organisms with species level IDs matched to IDs in the GlobDB database, which is a dereplicated set of species representative genomes [67]. The number of anti-phage defense systems per genome was determined using Defense Finder v2.0.1 [38]. To compare defense systems in non-bloom forming organisms, twenty random subsets of twenty genomes were extracted from GlobDB and annotated with Defense Finder. The number of defense systems per Mb of each random subset was compared to a random subset of twenty bloom forming organisms using a two-sided t-test.

We built a phylogenetic tree of bloom forming organisms by first building an alignment of a set of 120 ubiquitous single-copy genes (bac120) [68] with gtdbtk align v2.4.1 [46]. We used FastTree v2.1.3 [69] to construct a tree with the LG+CAT model with the *-gamma* parameter. We visualized the tree using iTOL [70].

## Results

### Virus-like particle (VLP) abundance decreased at high cell densities

TRL blooms are dominated by a species-level lineage of anoxygenic phototrophs within the *Prosthecochloris* genus: GSB-TRL01 [8]. In B01 in 2021, cell densities peaked at ∼6 × 10^7^ cells mL^-1^ two days after the onset of the bloom. Cell numbers then declined until the bloom disappeared (figure 2a, supplementary table 2). In B05, cell densities reached a peak of ∼10^9^ cells mL^-1^ six days after the start of sampling, increasing ∼10 fold over just 48 hours (figure 2a). This corresponds to a growth rate of ∼1.1 day^-1^ and a doubling time of ∼15 h. Typical cell densities in TRL outside of the bloom and after bloom termination were in the 10^6^ to 10^7^ cells mL^-1^ range (figure 2, last timepoint in b-d, supplementary figure S1a, and supplementary table 2). Fluorescence in situ hybridization revealed that nearly all cells (83.2-95.5% relative abundance) belonged to the Chlorobiota (figure 2a, supplementary table 2).

**Figure 2.**
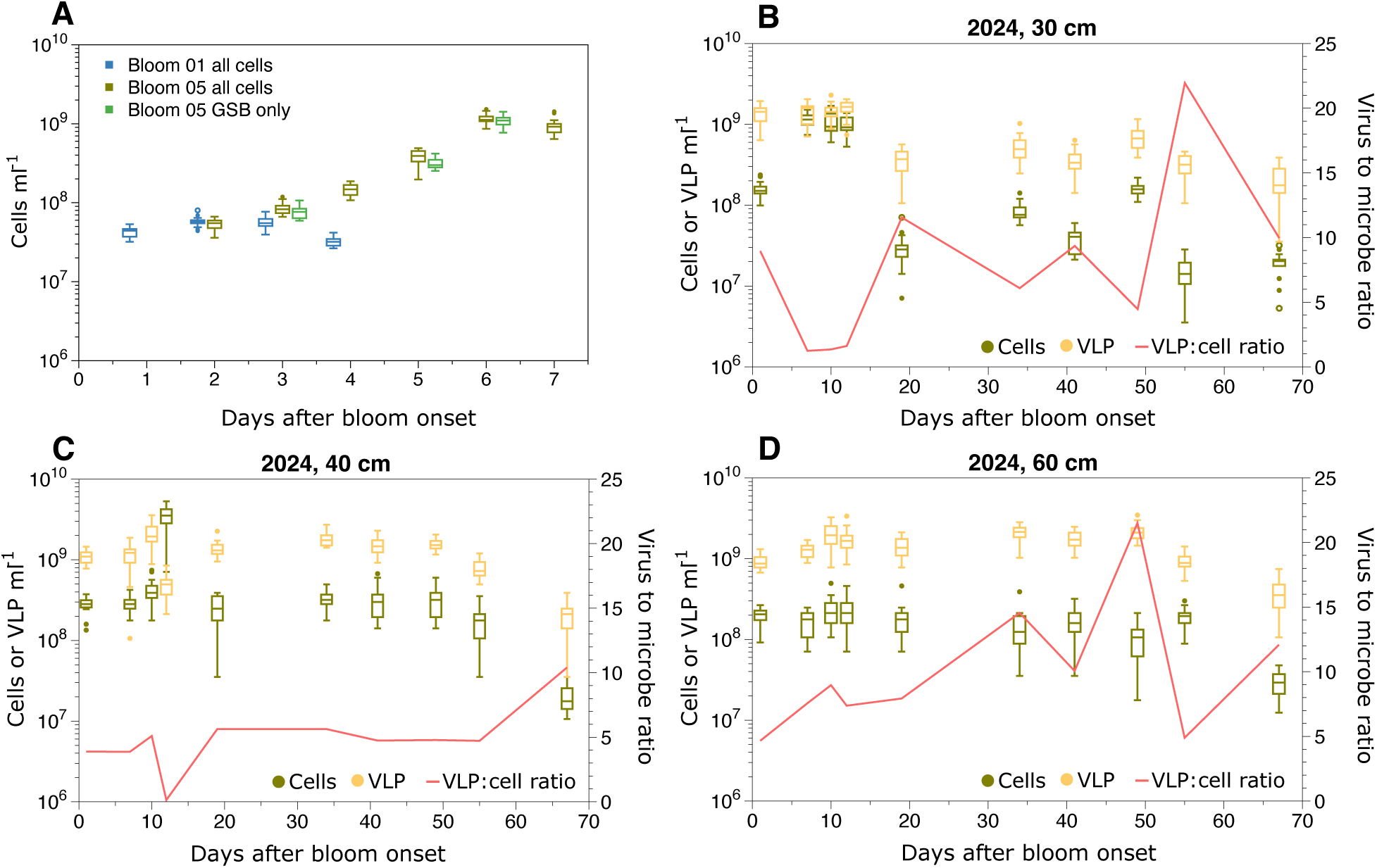
Total cell and virus-like particle (VLP) abundances and Chlorobi specific cell abundances at different timepoints throughout the bloom. (a) In 2021, total cell and VLP abundances are shown for two different blooms, B01 (blue) and B05 (brown green), GSB cell abundances were determined only for B05 (light green). (b-d) All total cell (brown green) and VLP abundances (yellow) are from different depths of bloom B05. The red line is the VLP to microbe ratio. For all samples 20 fields of view were counted (n=20 for each time point) summarized by standard boxplots.

In 2024, cell densities at 10 cm (above the bloom, S1a) and 65 cm (below the bloom, S1b) remained relatively constant throughout the time series. Cell densities within the bloom at 30 cm, 40 cm, and 60 cm (figure 2b-d) fluctuated over time as the bloom formed and eventually terminated. At 40 cm depth we found the highest cell densities (∼2 × 10^9^ cells mL^-1^ 12 days after the start of sampling), again increasing ∼10-fold over 48 h, corresponding to an approximate growth rate of ∼1.15 day^-1^ and a doubling time of ∼14.5 h (Figure 2b-d).

VLP abundances were relatively constant in the 10 cm and 65 cm layers. In these two layers, the VLP:cell ratio was consistently close to ten. In the 30, 40 and 60 cm layers VLP:cell ratios were more variable. When cell densities were highest, 7-12 days after the start of sampling, VLP:cell ratios dropped to ∼1. This reduction in VLP:cell ratios was particularly extreme at 40 cm twelve days after the start of sampling, when viral abundances were lower than cell abundances, causing the VLP:cell ratio to drop to ∼0.1.

### GSB-TRL01 is dominant during the bloom, and viral coverage patterns suggest active infection

Based on relative abundances from read mapping, GSB-TRL01 was very abundant during all sampling periods (figure 3a, supplementary table 3a). In 2021, GSB-TRL01 and a conjugative plasmid were present in roughly the same relative abundance (figure 3a). In 2024 and 2025 the plasmid was less abundant (roughly 15% to 30% of the relative abundance of the genome). The sequence-based relative abundance increased with cell density as measured by cell counts and peaked seven days after the start of sampling (figure 3a).

**Figure 3.**
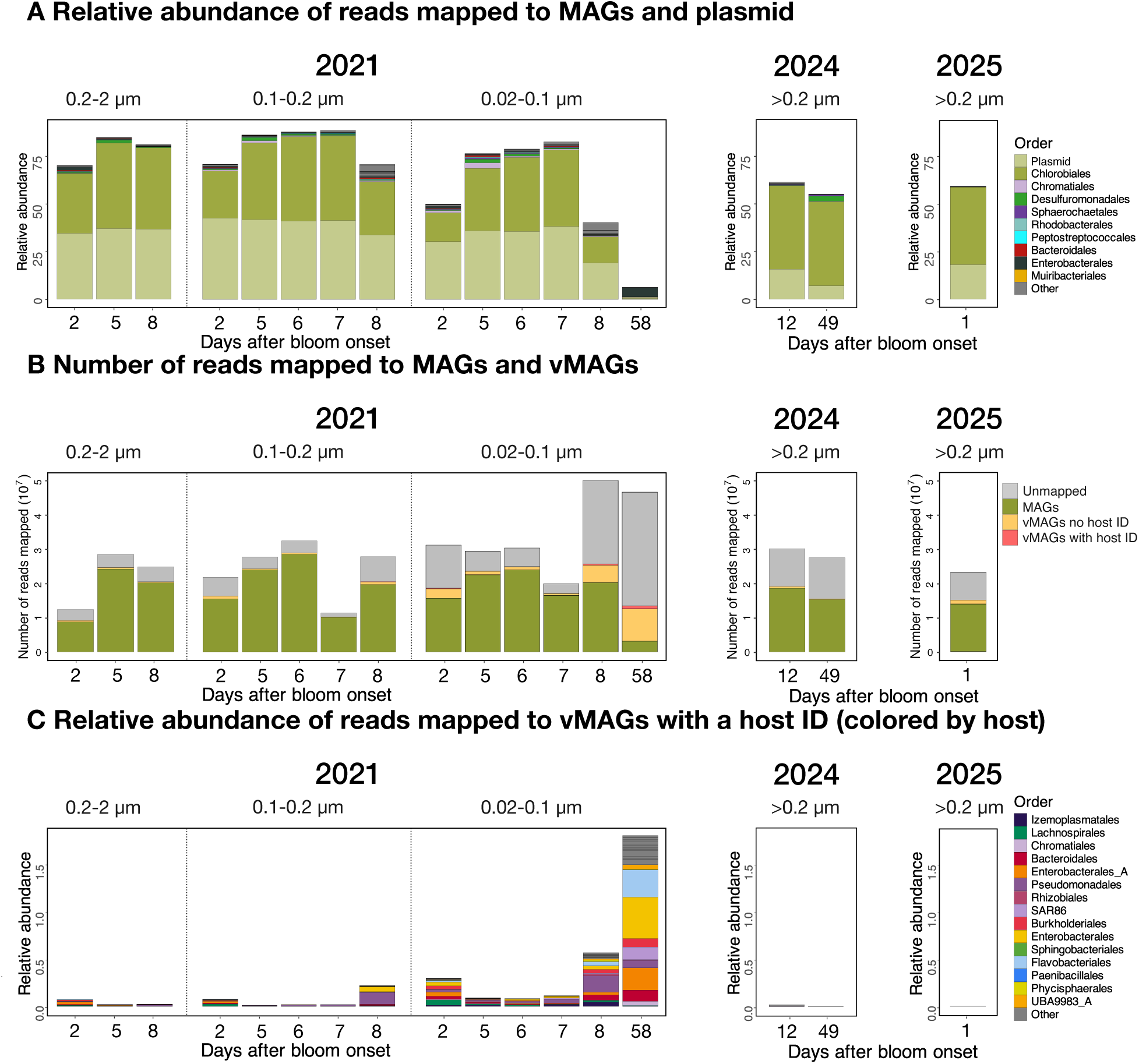
Relative abundances of MAGs, plasmid, and vMAGs over time series. a) Relative abundances of 95% ANI dereplicated MAGs and the conjugative plasmid. b) raw metagenomic read numbers mapped to MAGs (green), vMAGs without a host ID (yellow), vMAGs with a host ID (orange), and unmapped (gray). c) relative abundances of 97% ANI dereplicated vMAGs with a host ID, colored by the taxonomic identity of their putative host. Relative abundances were generated by mapping the Illumina short reads to MAGs and vMAGs and were calculated using the proportion of mean coverage rather than the proportion of reads to account for genome length. Relative abundance calculations are separated by the year in which samples were collected and were performed within each size fraction for samples from 2021. Note that c) represents host-matched viruses, which constitute the minority of viral reads matched, only.

Viral coverage generally decreased at the height of the bloom, possibly due to the increasing coverage of GSB-TRL01. There are, however, several vMAGs whose coverage increases during the bloom: vRhyme_bin_7, vRhyme_bin_1609, vRhyme_bin_3, and k141_71479 (all *Caudoviricetes* with no host ID, figure S3, supplementary table 3b). These four vMAGs had relatively high coverage in 2021 and 2024, but in 2025 only vRhyme_bin_1609 appeared with high coverage (figure S3). vRhyme_bin_1609 has a particularly high TPM across all three years. Despite the dominance of GSB-TRL01, viral metagenome assembled genomes (vMAGs) could not be bioinformatically linked to this organism using iPHoP (see supplement for discussion of limitations of this method). Although not matched to GSB-TRL01 with iPHoP, the vMAGs discussed above are candidates for successful infection during the bloom. A more detailed description of viral results is available in the supplement.

### The genome of GSB-TRL01 is enriched with defense mechanisms and associated with a conjugative plasmid

From Nanopore sequencing data, we recovered two apparently circular contigs with very high coverage. One contig was the complete 2.2 Mbp genome of the dominant bloom former in this system, GSB-TRL01 (Nanopore average read coverage = 150×, Illumina average read coverage = 834×, 99.99% completeness, 0.34% contamination).

The genome encoded 13 anti-mobile genetic element systems (supplementary table 4), including a Class 1 Subtype IC CRISPR-Cas system, five abortive infection systems (including the Thoeris system [71] and the recently characterized Gao_Ppl system [72]), four Type I restriction modification (RM) systems, a Wadjet anti-plasmid system, and two systems with demonstrated anti-phage defense activity, but undescribed mechanisms: HEC-03 [73] and VP1853 [74] (figure 4a). Wadjet, Gao_Ppl, VP1853, and an RM system clustered into a putative defense island on the genome (figure 4c). This area was enriched with transposases. The genome also contained a putative phage satellite (figure 4a, Supplementary figure S4, see supplementary results).

**Figure 4.**
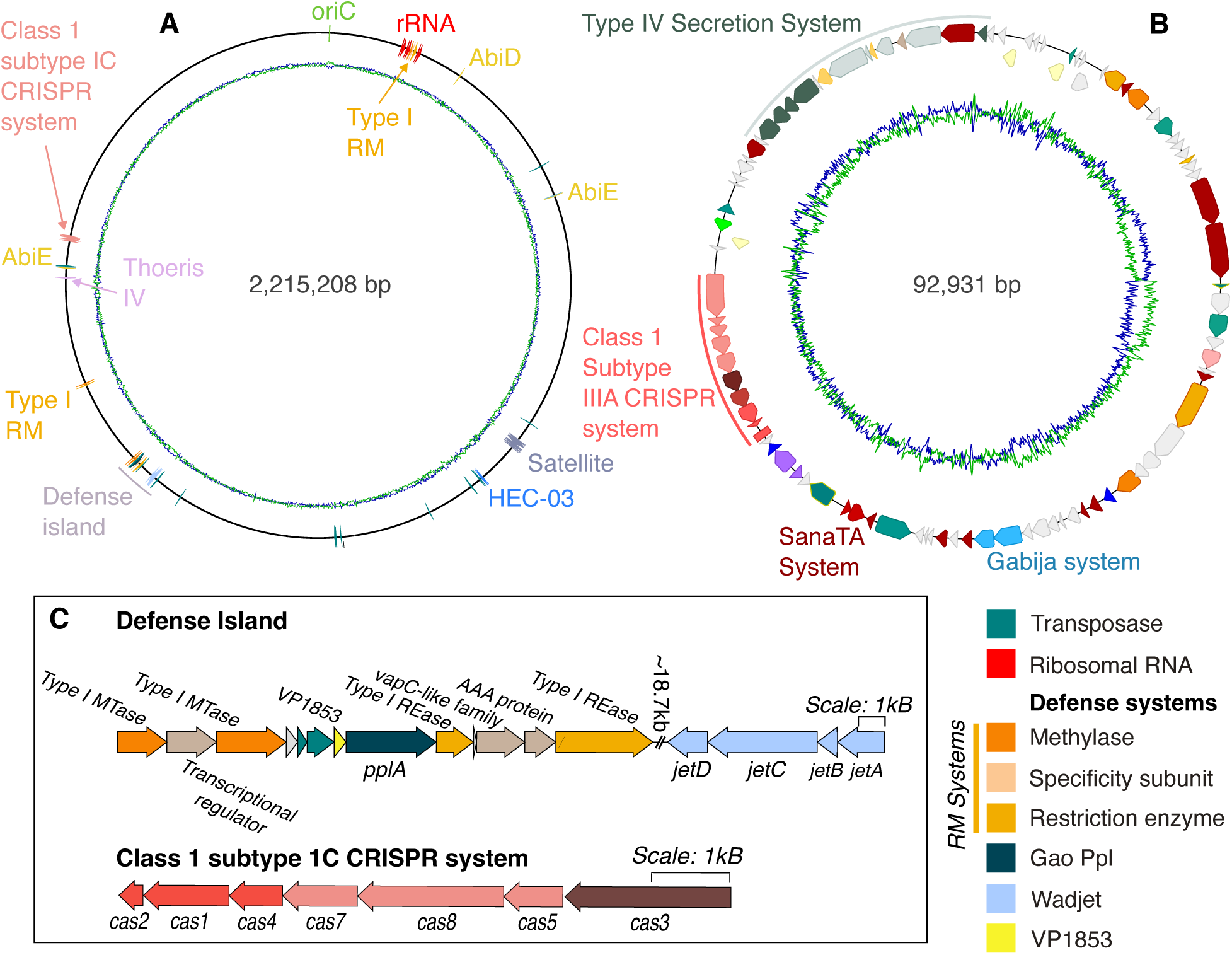
Map of the GSB-TRL01 genome and conjugative plasmid recovered from Nanopore sequencing. Both a) and b) represent singular contigs with contig length displayed in the center. In a) only genes related to mobile genetic elements, the oriC, and rRNA are displayed, whereas in b) all genes are represented. In both a) and b) coding sequences related to defense systems and mobile genetic elements are accompanied by a text label, or if unlabeled, color coded. Abi=abortive infection system, RM=restriction modification system. Green and blue lines represent AT/GC skew, respectively. c) Gene maps of the defense island and CRISPR system displayed on the GSB-TRL01 genome in a).

The other circular contig recovered from Nanopore sequencing was a 92 kbp conjugative plasmid, likely associated with GSB-TRL01 given its high coverage depth (Nanopore average read coverage = 167×, Illumina average read coverage = 1010×). Based on coverage, the plasmid was present at a ∼1.15:1 ratio with the genome, suggesting that most or all cells contained a copy, and some may have had multiple copies. Although an origin of transfer region was not identified, the plasmid contained a complete Type IV Secretion System and a MobP1 Relaxase (figure 4b), indicating that it was a conjugative plasmid. The plasmid additionally encoded a class I subtype IIIA CRISPR-Cas system, two Type II Restriction Modification systems, a SanaTA toxin-antitoxin system, and a Gabija abortive infection system (figure 4b, supplementary table 4).

### Expression of anti-phage defense proteins from the genome and the conjugative plasmid

We detected expression of genes from defense systems located on both the plasmid (4 systems) and the main genome of GSB-TRL01 (10 systems) in the metaproteomes, indicating that both are actively involved in defense against mobile genetic elements (figure 5, supplementary table 5). The relative abundance of Cas proteins was similar for proteins that are part of the CRISPR-Cas system in the main genome and on the plasmid. Average relative abundance of proteins from both CRISPR-Cas systems increased later in the bloom in both B05 (figure 5c) and B01 (figure S5). Genes from the Wadjet system, Gao_Ppl, six restriction-modification systems, the Gabija system, HEC-03, and toxins from two abortive infection systems were also expressed (figure 5, supplementary table 5).

**Figure 5.**
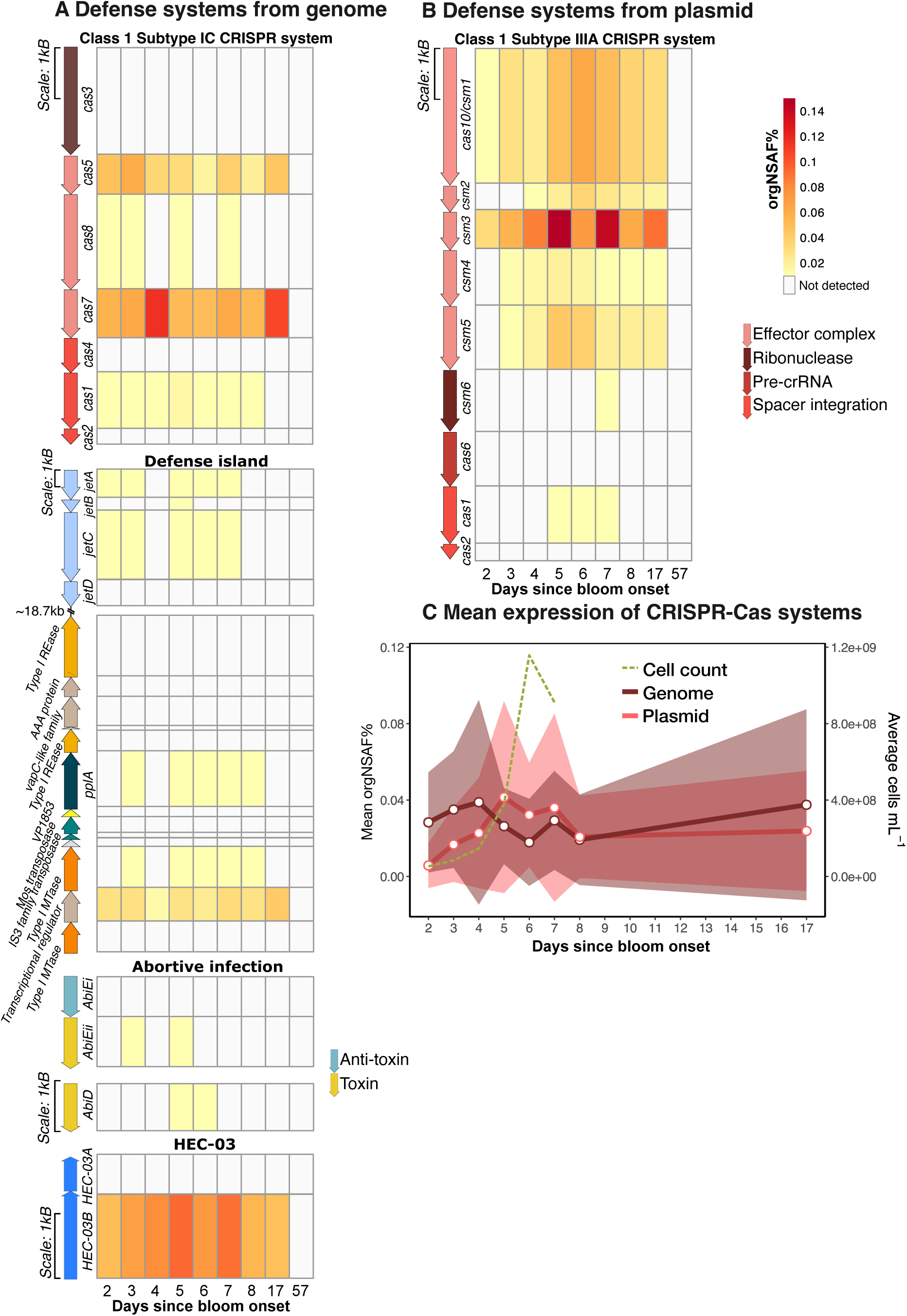
Expression of defense system genes. a) shows proteins of the cellular fraction of the GSB-TRL01 genome and b) of the associated plasmid. Heat maps show the protein abundances as organism normalized spectral abundance factor percent (orgNSAF%) in the B05 samples (see figure S5 for B01 values). In addition to the defense systems displayed here, proteins from four additional RM systems and the Gabija system were expressed (supplementary table 5). c) shows the mean (solid line) and standard deviation (shading) orgNSAF% for CRISPR-Cas systems in the genome (purple) and plasmid (pink) and average total cell counts for each time point (green, dashed). All orgNSAF% values have been normalized to reflect within-organism relative abundance only, meaning that the NSAF% value for each protein has been divided by the sum of all NSAF% values for each protein in the genome and the plasmid for each sample.

Genes throughout the plasmid were expressed. We detected proteins from the conjugative machinery of the Type IV Secretion System across most time points in both B05 and B01 samples (supplementary figure S6). In B05, T4SS proteins were more abundant in the early and late bloom stages, suggesting periods of more active plasmid transfer (supplementary figure S6). In B01, T4SS proteins were more abundant later in the bloom except the virB9 protein, which showed high relative abundance throughout the sampling period (supplementary figure S6).

### Detection of viral structural proteins suggests active infection

Structural genes from several viruses were more abundant in the proteome during the height of the bloom (supplementary figure S7, supplementary table 6). These phages, although they could not be bioinformatically linked to the Chlorobiota, are nonetheless candidates for GSB-TRL01 infecting phages. A major capsid protein from vRhyme_bin_7 (classified by geNomad as belonging to the *Kyanoviridae*, a family of *Caudoviricetes* of the Myovirus morphotype of which all known representatives infect Cyanobacteria [75]) was present in both the cellular and extracellular samples from the proteome. There are several other viruses which are GSB-TRL01 phage candidates based on increased extracellular proteomic abundances at the height of the bloom: k141_1026370, k141_127641, vRhyme_bin_38, k141_85354, vRhyme_bin_633 and others (supplementary figure S7). The absence of proteins from these phages in the cellular fraction may reflect short infection timescales which were not captured by once-daily sampling, or that the extracellular phages were generated by successful infections in only a very small fraction of the cellular community which was not captured in the cellular sampling.

To assess viral contribution to total biomass, we compared the peptide spectrum matches (PSMs) for viral proteins to all detected non-viral proteins (supplementary figure S8, supplementary table 7). In the cellular metaproteomic samples the viral contribution to total biomass ranged from ∼0.15% to ∼0.85% in B05 and ∼0.17% to ∼0.29% in B01. In the extracellular metaproteomic samples the viral contribution to total biomass ranged from ∼0.5% to ∼12% in B05 and ∼0.2% to ∼1.4% in B01. In both blooms, the viral percent contribution to total biomass decreased during the highest cell densities as compared to pre and post bloom timepoints. While extracellular viral biomass percentages are mostly below 3%, 57 days after the start of the bloom in B05 there was a striking peak of ∼12% viral biomass.

### Defense systems are enriched in numerous bloom-forming prokaryotes affiliating with diverse phylogenetic lineages

To understand whether increased investment in phage defense is a general trait in prokaryotic bloom-formers, we carried out a large-scale meta-analysis. We collected and analyzed 89 species of putative bloom formers (four Archaea and 85 Bacteria) from studies conducted in diverse aquatic environments (supplementary table S8). We identified the species representative for each prokaryotic bloom former in GlobDB [67] (a dereplicated database of species representative genomes and MAGs), then compared random subsets of these bloom formers to random subsets of putative non-bloom forming organisms. In this comparison, bloom forming organisms had significantly more defense systems (figure 6a). The average number of defense systems per Mb among random subsets of twenty bloom forming organisms was ∼4.5 systems, whereas for random subsets of non-bloom forming organisms, the average was ∼1.5 systems. In twenty t-tests of random subsets of twenty bloom forming organisms compared to twenty non-bloom forming organisms, the p-value was <0.05 (figure 6a).

**Figure 6.**
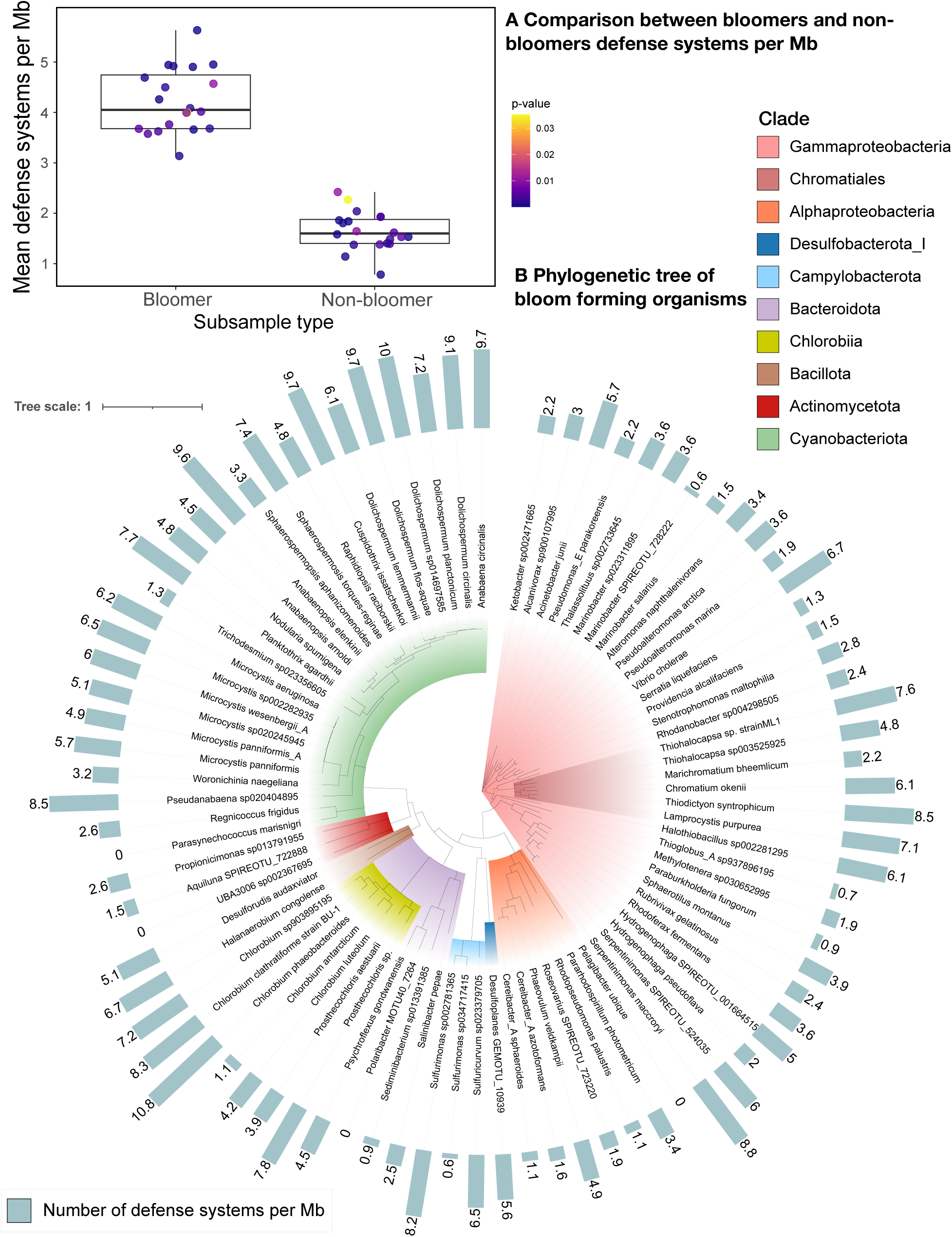
Diverse bloom-forming bacteria encode abundant defense systems. a) A comparison of the number of defense systems per mega-base (Mb) between bloom forming organisms and non-bloom forming organisms. Each point represents the mean number of defense systems per Mb of a randomly selected group of twenty organisms. The color of the point represents the p-value of a t-test conducted between the two randomly selected groups (this comparison was repeated 20 times). b) A phylogenetic tree of bacteria identified as bloom-forming in this study. The outer bar graph represents the number of defense systems per Mb for each organism. Colored regions represent clades.

The number of defense systems varied by taxonomic identity. Three clades were particularly enriched: the phylum *Cyanobacteriota*, the class *Chlorobiia*, and the order *Chromatiales* (figure 6b). Archaea (n = 4, average 7) (not shown in figure 6b) tended to have lower numbers of defense systems than did Bacteria (n = 85, average 19).

## Discussion

### Cell and VLP abundance dynamics indicate lower rates of lytic infection at high cell densities

The VLP:cell ratio was approximately 10:1 at low to moderate cell densities. However, at the highest cell densities, this ratio dropped substantially (with a minimum at ∼0.1 VLP:cell ratio at peak cell density) (figure 2). These data are supported by the observation that the viral percent biomass during peak cell densities was lower than percent biomass in samples from early and late in the bloom (supplementary figure S8). If GSB-TRL01 experiences rampant lytic infection, VLP abundance should increase with cell abundance. Instead, we see an opposite trend (figure 2). This contradicts the traditional “boom and bust” view of bloom dynamics, wherein lytic infection leads to an increase in free viral particles at high cell abundances, eventually leading to infection mediated bloom collapse. While virally mediated collapse may be a factor in the termination of some blooms [11–15, 76, 77], the pattern we observe, of reduced abundance of free VLPs at very high cell densities, has been previously observed in anoxygenic phototrophic blooms [1, 3, 16]. Wigington et al. find that viral abundances have a non-linear relationship with cellular abundances across marine environments, fit by a power law relationship in which the VLP:cell ratio decreases with increasing cell density [78]. The VLP to cellular abundance relationship in our direct count data is fit by a power law, except at the highest cellular abundances (figure S9), suggesting that at very high cell abundances, there may be a shift in how VLP concentrations scale.

One possible explanation for this dynamic is that blooming organisms invest heavily in anti-viral defense, leading to lower rates of infection overall when blooming organisms are extremely dominant. This scenario is consistent with models like Kill-the-Winner [20] and density dependent fluctuating selection [79], wherein “defense specialists” can outcompete “competition specialists” under favorable conditions. While this model has been previously proposed to account for the high abundances of SAR11 cells [80] we are unaware of it having been explicitly tested in the context of bacterial blooms.

While the abundance of free viral particles declined at high cell densities, we provide evidence that infection likely occurs in the dominant bloom former. One vMAG in particular, vRhyme_bin_7, is a strong candidate for active infection given both metaproteomic and metagenomic evidence (see supplement for detailed discussion).

### GSB-TRL01 invests heavily in anti-phage defense mechanisms

The genome of GSB-TRL01 is enriched with 13 defense systems as compared to the average microbial genome (five systems) [81]. We provide evidence that each GSB-TRL01 cell additionally contains a conjugative plasmid, which encodes another five defense systems. The question of why certain bacteria accumulate more abundant and diverse defense systems than others is an emerging topic in the field of viral ecology [82]. Two important assumptions are commonly held in defense ecology and have been the subject of much investigation and discussion: a) that more defense systems mean better defense, and b) that defense systems are costly to the host. Recent work with bacterial culture collections and phages isolated from different environments did not demonstrate that increased abundance of anti-phage defense mechanisms led to increased defense ability [83]. However, studies that used hosts and phages from the same environment, a situation more reflective of ecological reality, did find a positive correlation between number of defense systems and defense ability [84, 85]. An additional survey of global metagenomic data found a positive correlation between phage abundance and number of defense systems encoded in microbial genomes, reinforcing the idea that additional defense systems provide stronger protection [66]. Given these results, we suggest that the accumulation of anti-phage defense systems in the GSB-TRL01 genome reflects its robust defense abilities and results from predation pressure in blooms.

Given that not all bacteria encode all defense systems, we assume that there is a cost of resistance (COR). The COR arises from multiple factors including metabolic costs, autoimmunity, and limiting horizontal gene transfer. These costs can result in the loss of defense systems that do not provide a sufficient benefit to the host. Additionally, hosts manage COR by carefully regulating the expression of defense systems [86]. In this section, we detail several examples of the potential costs of the 13 defense systems found in GSB-TRL01. The 13 defense systems found in the genome of GSB-TRL01 likely exert high costs on the host, both in terms of energy expenditure and autoimmunity risks. The acquisition and maintenance of a large conjugative plasmid is also costly for the host. Refer to the supplementary discussion for full discussion of the costs associated with defense systems and conjugative plasmids. The maintenance and expression of such a large repertoire of anti-phage defense systems therefore suggests a benefit in the form of phage resistance that outweighs the potential fitness costs of these systems. Similarly, the presence of a large conjugative plasmid, which was apparently present in every cell, may indicate that the host was deriving a substantial fitness benefit. As a photoautotroph, GSB-TRL01 has access to an abundance of energy, which may prevent energy-related costs of these systems from imposing a significant burden.

### Investment in defense may be a conserved trait among bloom forming organisms

Here, we demonstrate that bloom forming prokaryotes across the tree of life appear to be enriched with defense systems, supporting our hypothesis that bloomers are defense specialists. Interestingly, this enrichment seems to be exaggerated among phototrophic bloomers in the phylum *Cyanobacteriota*, the class *Chlorobiia*, and the order *Chromatiales* (figure 6b). The enrichment of defense machinery in bloom-forming Cyanobacteria has been previously observed [24–27]. This may suggest that defense is particularly important for phototrophic bloom formers, or that, as photoautotrophs with a potentially unlimited energy supply, they are particularly energetically suited to cope with the costs of additional defense systems.

Recently, there has been much interest in the question of why some organisms encode more defense machinery than others [82]. While recent papers have focused on the impact of environment and host/phage density on the prevalence of defense systems [66, 87, 88], fewer have considered host ecological strategy as a factor. Our analysis strongly suggests that considering ecological strategies is essential in the quest to understand why certain organisms have more defense systems than others.

Bloom-forming organisms are generally thought to be opportunistic organisms that grow to dominate under advantageous nutrient or environmental conditions. They are likely to have a fast growth rate under optimal conditions, have a high nutrient demand and low substrate use efficiency, and variable population dynamics. However, few studies have sought to experimentally generalize the qualities that make bloom formers successful. Here, we show that many have the genomic capabilities to be defense specialists, as outlined by Thingstad in the Kill-the-Winner hypothesis [19].

### Conclusion

We show that GSB-TRL01’s genome is enriched with a high number of anti-phage defense systems, and that it is associated with a conjugative plasmid carrying additional defense systems. Given the observed decrease in VLP counts at high cell density, we propose that heavy enrichment of anti-phage defense machinery helps these hosts suppress the rates of viral lysis, causing viral production to decrease when GSB-TRL01 is dominant. This investment in defense seems to be shared across most bloom forming organisms, further supporting the hypothesis that it is a mechanism for long-term bloom persistence. Bacterial blooms are important drivers of ecosystemic carbon flow, either by fixing carbon (in the case of phototrophic blooms), or remineralizing carbon (in the case of the heterotrophic bacterial blooms that often follow marine phytoplankton blooms). Our results show that anti-phage defense may play an important role in this carbon flow by determining the duration and rate of decline of bacterial blooms. Our findings may also be consequential for efforts to engineer microorganisms for industrial purposes, such as the production of biopolymers and biofuels. Understanding the ecological properties of successful bloom formers will be an essential piece of manufacturing phage-resistant synthetic “blooms” in industrial settings. Additionally, this study may be helpful for the development of phage therapy. Several of the organisms recovered in our bloom forming database are common pathogens, which have ecological strategies very similar to environmental bloom formers. Further investigation is necessary to clarify whether this trend of anti-phage enrichment applies broadly to pathogens, potentially explaining the differences in their responsiveness to phage therapy. Collectively, our findings illuminate a mechanism for survival and persistence in blooms, and will contribute to future efforts to understand and engineer the microbial world.

## Data Availability

Metaproteomic data, raw mass spectrometry files, and databases used are deposited in the ProteomeXchange Consortium via the PRIDE partner repository under the project accession PXD064785 [https://www.ebi.ac.uk/pride/login, token: ylrzohYwCArL] [89]. Metagenomic reads, MAGs, and vMAGs have been deposited in the NCBI BioProject PRJNA1417754. The additional files required to reproduce our analyses are available in an Open Science Framework data repository (https://osf.io/skuq7/overview?view_only=4ecb92165508457e8fdad89780139af3). Scripts used in analysis and to generate figures are available at https://github.com/moyn413/TrunkRiverChlorobi.

## Supplementary files

Supplementary table 1: Summary of all samples used in this study

Supplementary table 2: Summary of cell, VLP, and Chlorobi specific counts from B05 and B01 in 2021 and 2024

Supplementary table 3: Summary of MAG (a) and vMAG (b) quality, taxonomy, and relative abundances

Supplementary table 4: DefenseFinder annotations for GSB-TRL01 and conjugative plasmid.

Supplementary table 5: GSB-TRL01 and conjugative plasmid annotations and metaproteomic orgNSAF% expression values for proteins represented in the metaproteome.

Supplementary table 6: Annotations and metaproteomic NSAF% values of vMAG proteins represented in the metaproteome

Supplementary table 7: Calculated percent contribution to biomass across samples for each vMAG

Supplementary table 8: Database of bloom forming organisms

## Acknowledgements

LC-MS/MS measurements were carried out in the Molecular Education, Technology, and Research Innovation Center (METRIC) at North Carolina State University, which is supported by the State of North Carolina, USA. Bioinformatic analyses were carried out at the JBPC Keck Computational Biology Lab. We thank Richard Fox for excellent bioinformatics support and Sandra Kolundžija for her help developing our VLP counting protocol.

