## Supplementary text for "Bloom-forming bacteria heavily invest in anti-phage defense"

[1] The Marine Biological Laboratory, Woods Hole, MA, 02543, USA
[2] The University of Chicago, Chicago, IL, 60637, USA
[3] Florida Atlantic University, Boca Raton, FL, 33431, USA
[4] North Carolina State University, Raleigh, NC, 27695, USA
[5] UC Santa Barbara, Santa Barbara, CA, 93106, USA
[6] Virginia Polytechnic Institute and State University, Blacksburg, 24061, USA
[#]present addresses:
EG: University of Amsterdam, Netherlands
CAC: University of Connecticut, Groton, CT, USA
ENJ: Nantucket Conservation Foundation, Nantucket, MA, 02554, USA

#### Supplementary figures

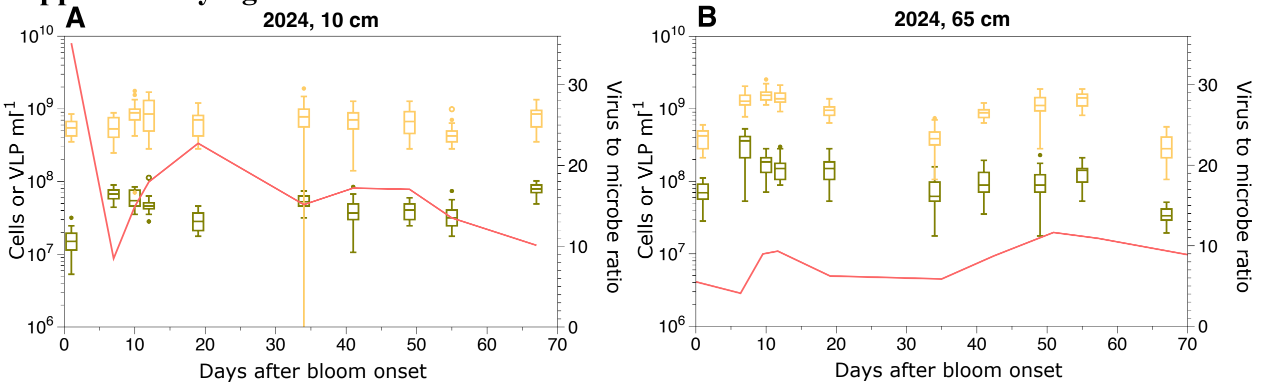


**S1. Direct cell and virus-like particle (VLP) counts**. Counts were conducted with epifluorescence microscopy (with DAPI and Sybr Green I respectively) from the 10 cm depth (a) and the 65 cm depth (b). Cell counts are total counts from B05 (asparagus green). Total VLP counts are from the same samples as total cell counts, but pre-filtered with a 0.2 µm Amicon filter (yellow). The orange line is the VLP:cell ratio. For all samples 20 fields of view were counted (n=20 for each time point).

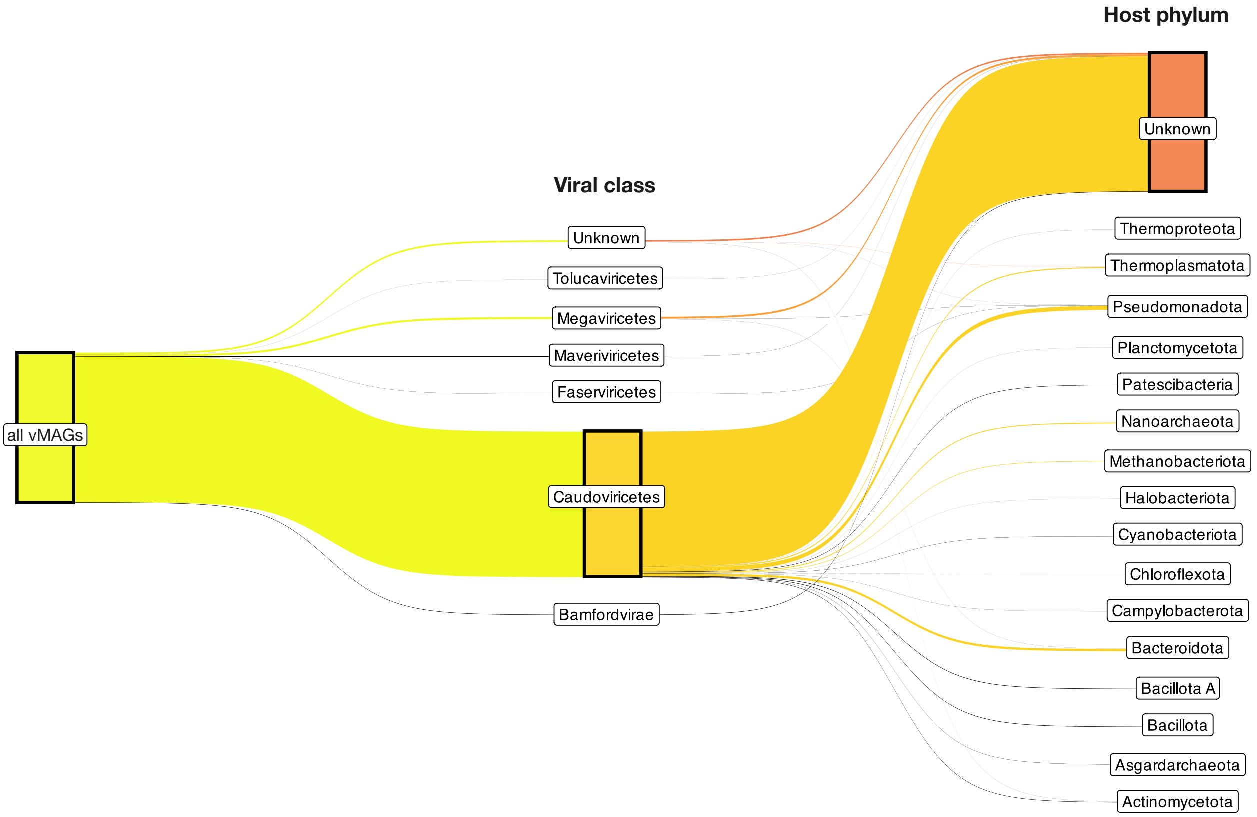


**S2. Sankey diagram showing the fractionation of total vMAGs into viral class (as classified by geNomad and ViralRecall), then into host identification (as identified by iPHoP).**

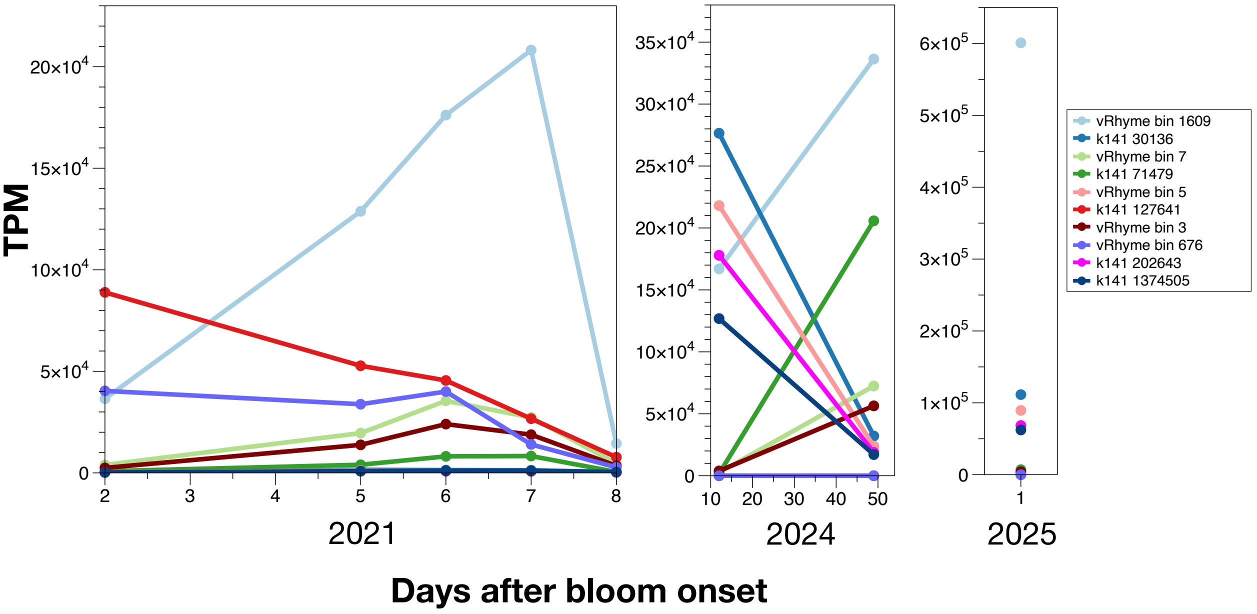


**S3. Reads per kilobase million (TPM) mapped to the 10 vMAGs with the highest total TPMs across the three sampling years.** Coverages come from the reads from the 0.025-0.1 µm size fraction in 2021, and the >0.2 µm size fraction in 2024 and 2025. Note that the y-axis has a different scale for each year.

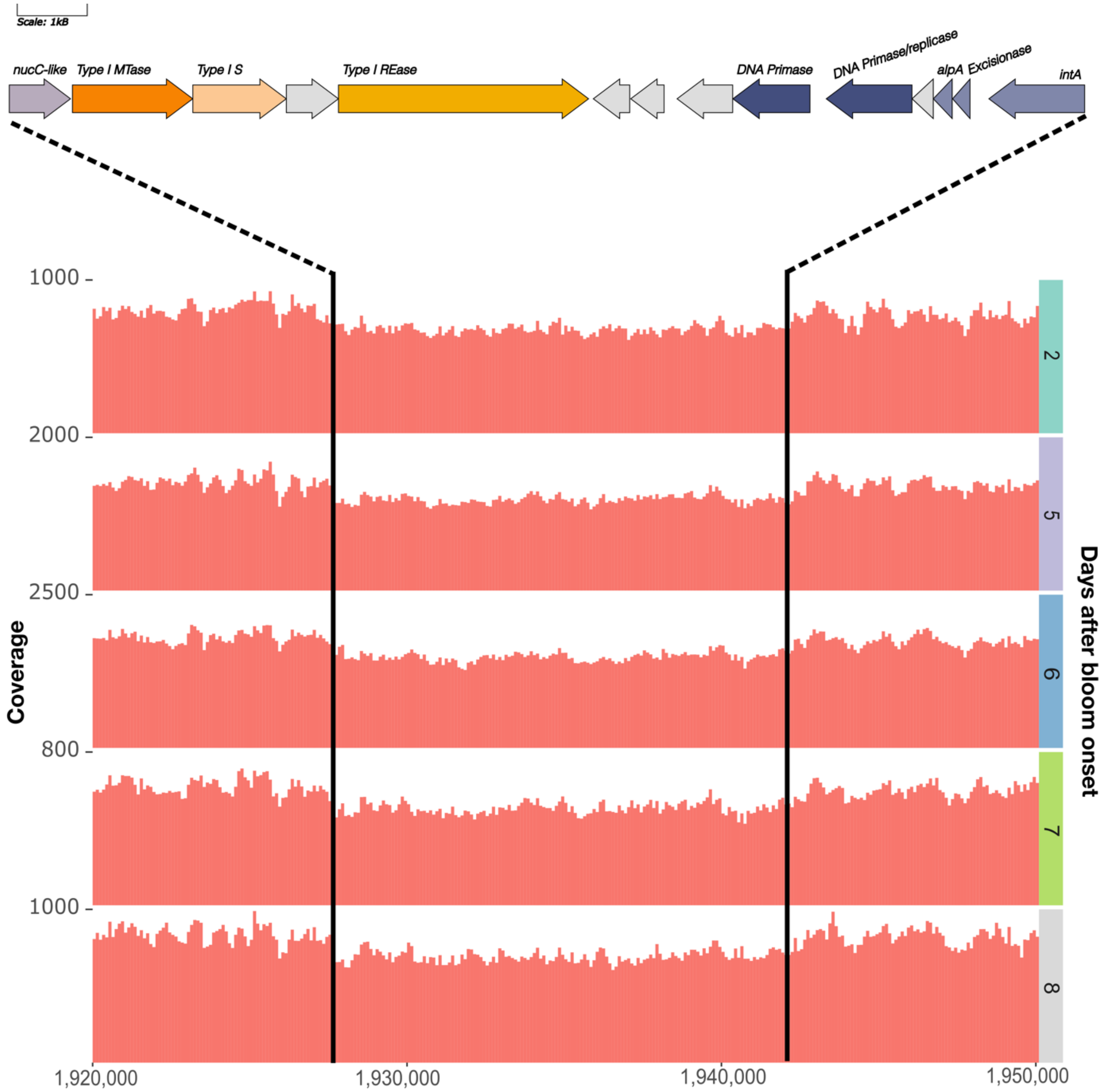


**S4: Gene map of putative phage inducible chromosomal island (PICI) Type C variant 1 satellite (~14kb long) and Illumina short read coverage over the 2021 time series.** Shades of orange represent a complete Type I RM system. The purple gene on the left end of the gene map is a nucC-like gene and may represent a fragment of a CBASS system. Shades of blue represent key functional genes. Grey genes are unannotated. All coverage plots are from the 0.1 µm to 0.2 µm size fraction and plots are faceted by time point. The area between the black lines corresponds to the PICI.

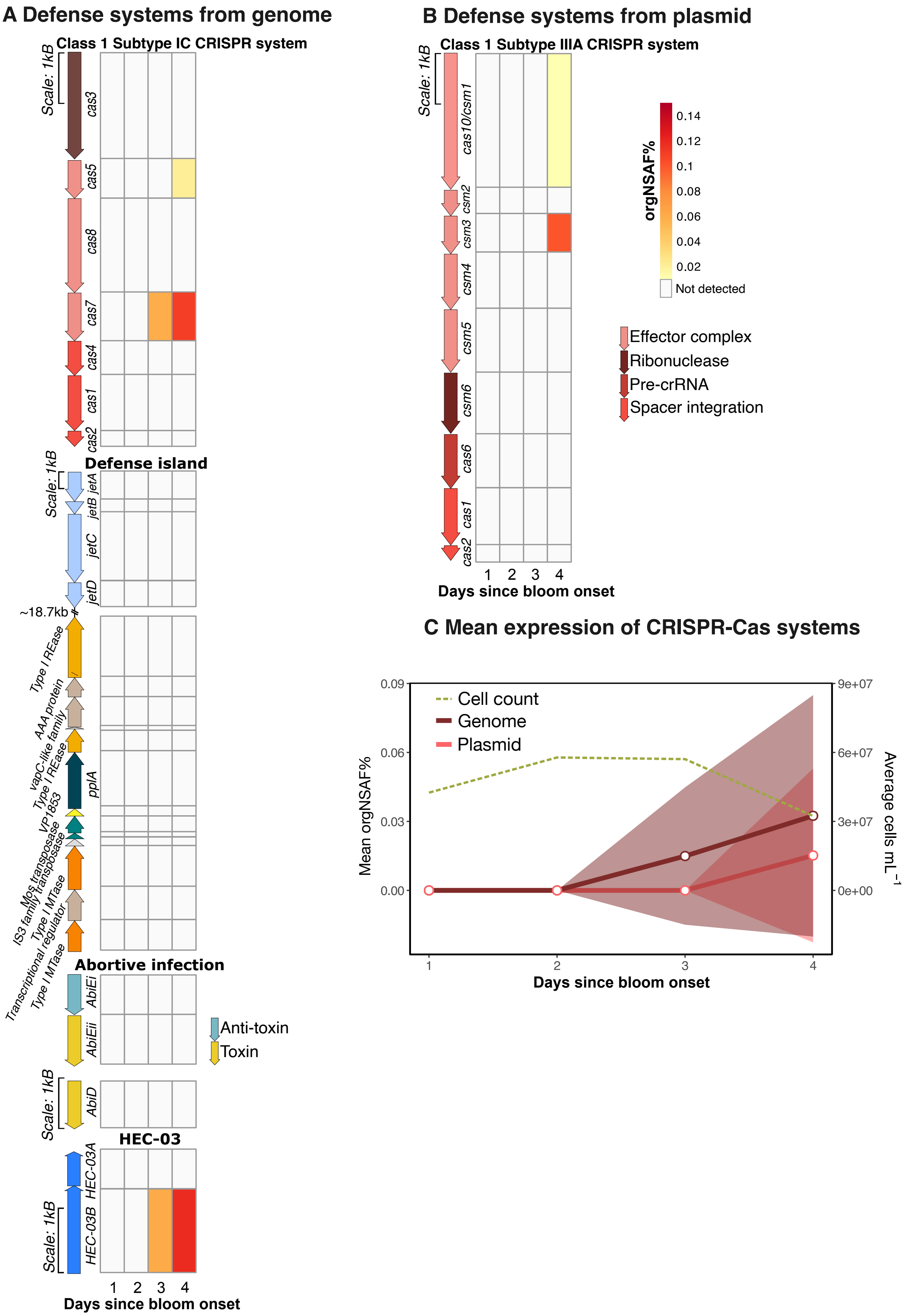
A Defense systems from genome
Class 1 Subtype IC CRISPR system
Scale: 1kB
cas3
cas5
cas8
cas7
cas4
cas1
cas2
Defense island
Scale: 1kB
jetA
jetB
jetC
jetD
~18.7kb
Type I REase
AAA protein
vapC-like family
Type I REase
pplA
VP1853
Mos transposase
IS3 family transposase
Type I MTase
Transcriptional regulator
Type I MTase
Abortive infection
AbiEi
AbiEii
Anti-toxin
Toxin
Scale: 1kB
AbiD
HEC-03
HEC-03A
HEC-03B
Scale: 1kB
1 2 3 4
Days since bloom onset
B Defense systems from plasmid
Class 1 Subtype IIIA CRISPR system
Scale: 1kB
cas10/csm1
csm2
csm3
csm4
csm5
csm6
cas6
cas1
cas2
1 2 3 4
Days since bloom onset
orgNSAF%
0.14
0.12
0.1
0.08
0.06
0.04
0.02
Not detected
Effector complex
Ribonuclease
Pre-crRNA
Spacer integration
C Mean expression of CRISPR-Cas systems
Cell count
Genome
Plasmid
Mean orgNSAF%
0.09
0.06
0.03
0.00
9e+07
6e+07
3e+07
0e+00
Average cells mL$^{-1}$
1
2
3
4
Days since bloom onset

**S5. Expression of defense system genes measured by metaproteomics**. Expression in the cellular metaproteome from a) the GSB-TRL01 genome and b) the associated plasmid. Heat maps show the protein abundances as normalized spectral abundance factor percent (orgNSAF%) in the B01 samples (see figure 5 for B05 values). Only defense systems with proteins detected in the cellular metaproteome are displayed, but not all systems with expression are shown. c) shows the mean (solid line) and standard deviation (shading) orgNSAF% for CRISPR-Cas systems in the genome (purple) and plasmid (pink) and average total cell counts for each time point (green, dashed). All orgNSAF% values have been normalized to reflect within-organism relative abundance only.

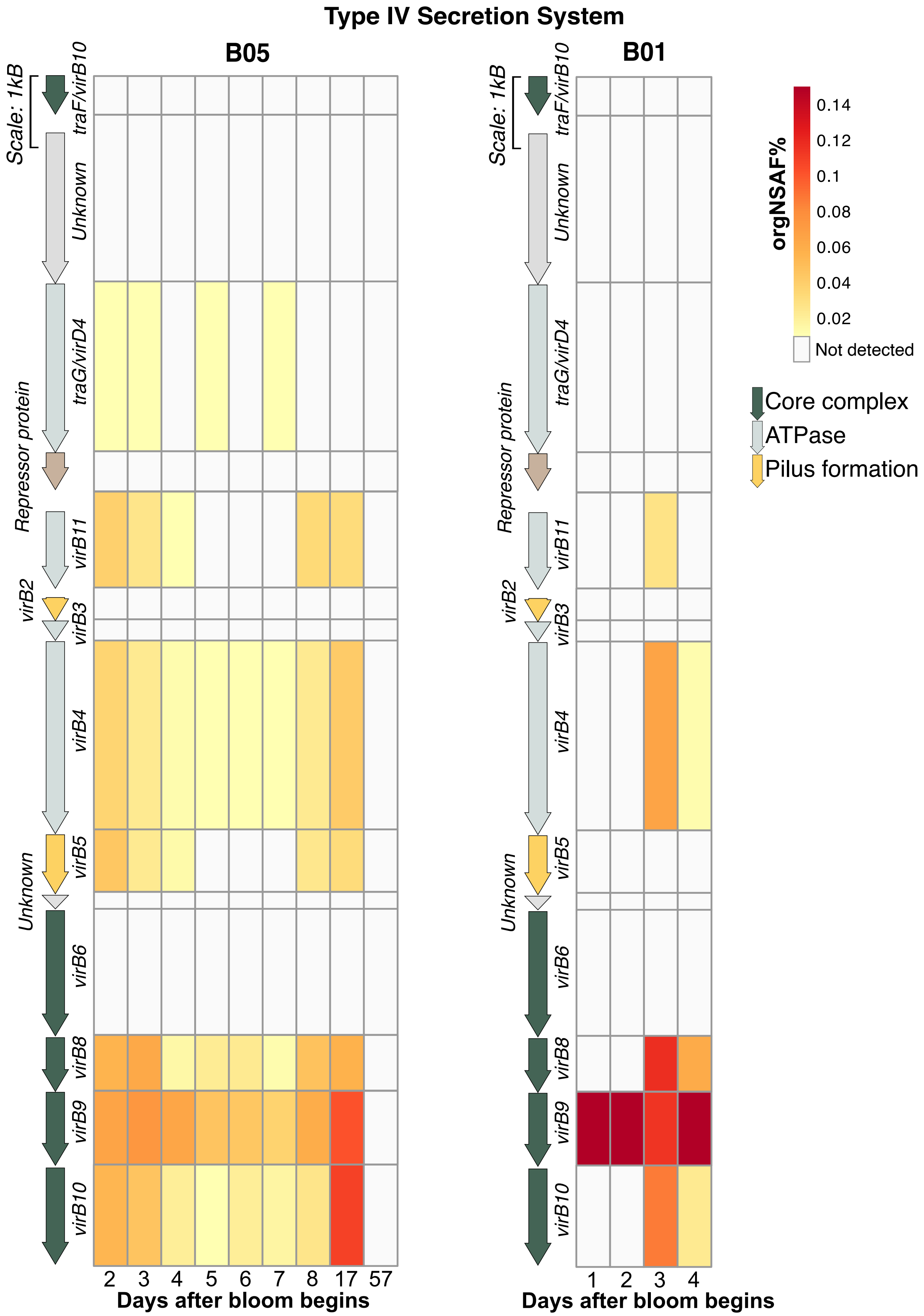

Type IV Secretion System
B05
B01
Scale: 1kB
traF/virB10
Unknown
traG/virD4
Repressor protein
virB11
virB2
virB3
virB4
virB5
Unknown
virB6
virB8
virB9
virB10
2 3 4 5 6 7 8 17 57
Days after bloom begins
1 2 3 4
Days after bloom begins
orgNSAF%
0.14
0.12
0.1
0.08
0.06
0.04
0.02
Not detected
Core complex
ATPase
Pilus formation

**S6. Expression of the plasmid's Type IV secretion system as measured by metaproteomics.** Gene map of the plasmid's Type IV secretion system, accompanied by a heat map showing the proteomic percent normalized spectral abundance factor (orgNSAF%) for each gene in B05 (left) and B01 (right) cellular metaproteomic samples.

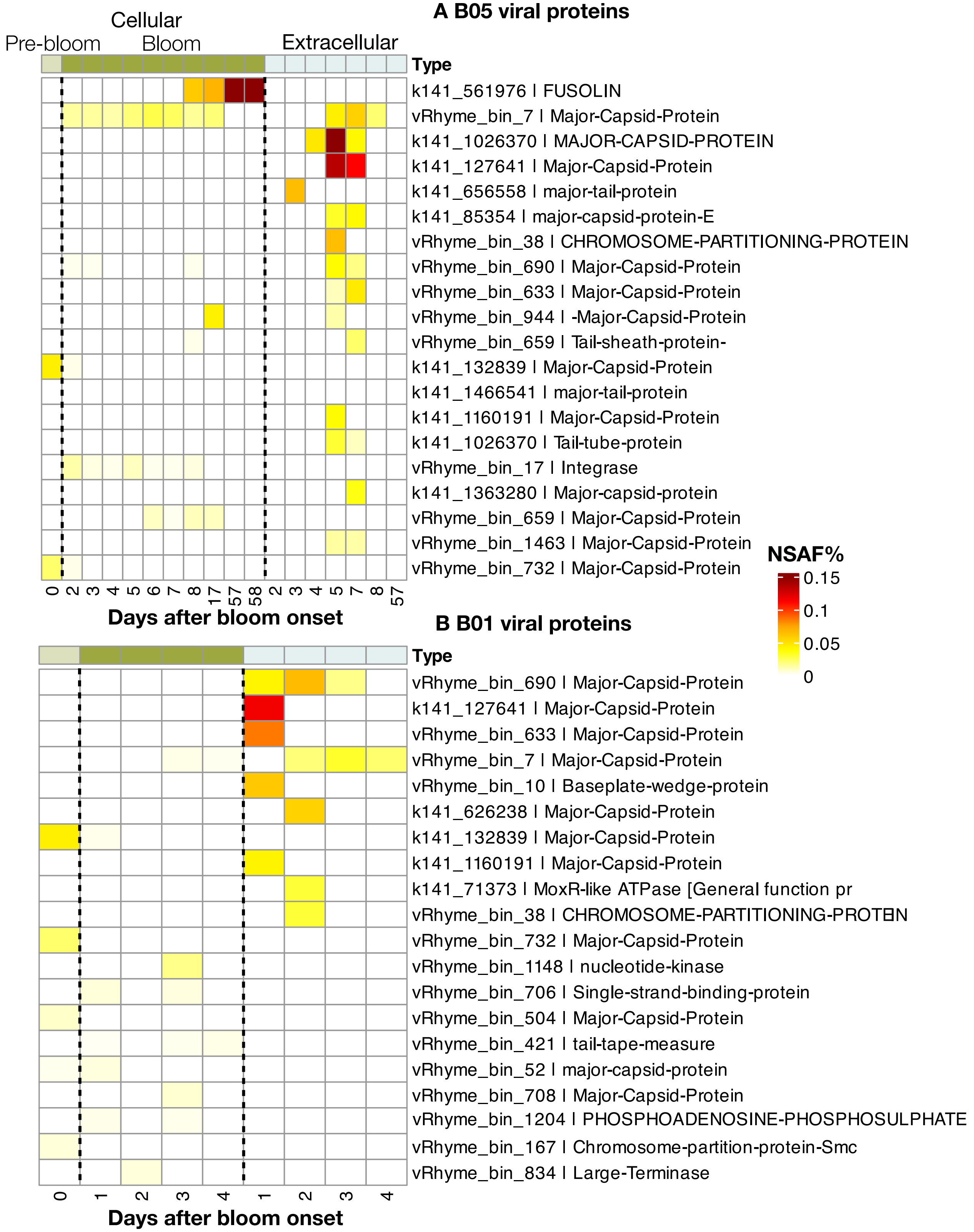


**S7. Viral proteomic percent normalized spectral abundance factor (NSAF%) expression over the course of the bloom for the top 20 most abundant viral proteins in a) B05 and b) B01.** The x-axis represents days after the start of the bloom and is subdivided into three sections.

The leftmost two fractions represent the intracellular fraction from the pre-bloom (light green) and bloom (asparagus green). The pre-bloom sample is the same in a) and b). The fraction on the right is extracellular. Each row of the heatmap represents one protein. The names of the proteins are represented by “name_of_vMAG : description of gene”, where the description of gene is as-given by Cenote-Taker3.

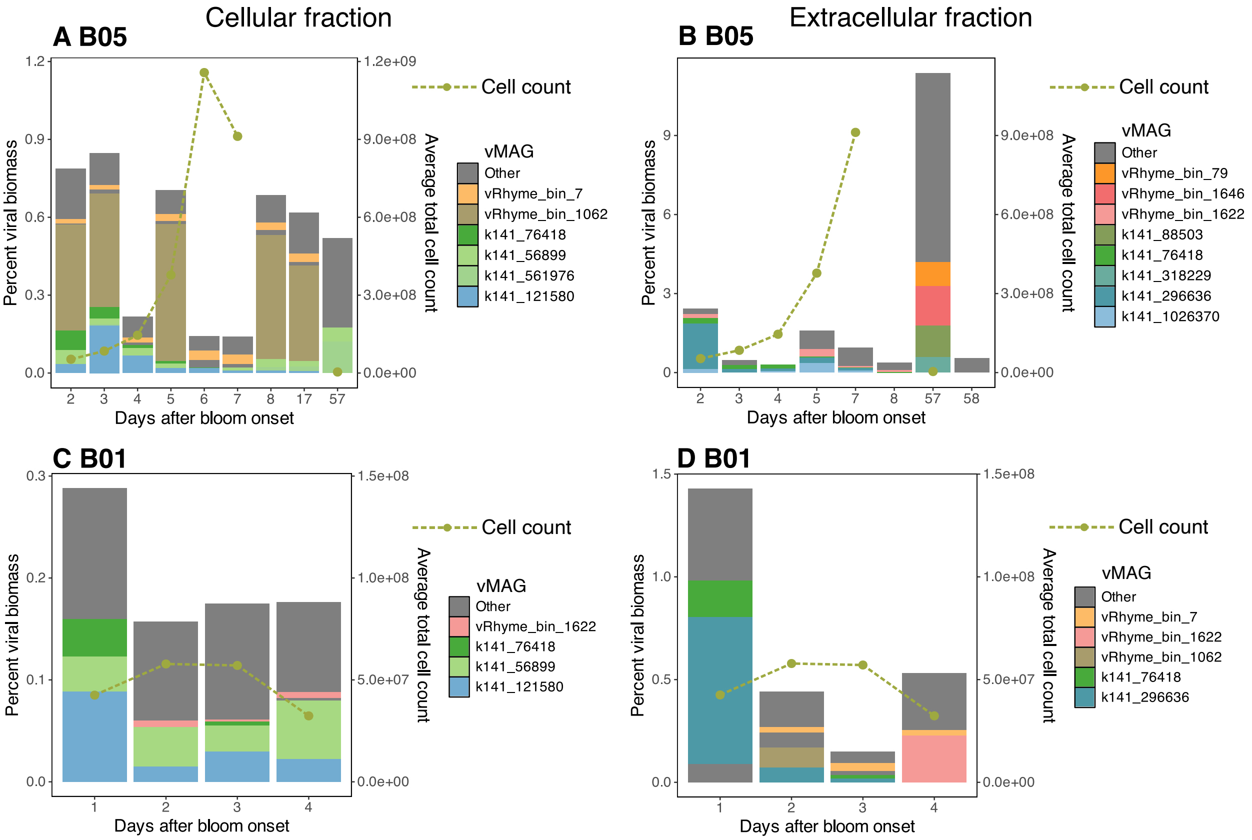


**S8. Viral percent contribution to total biomass in the metaproteome.** Percent biomass calculated by dividing peptide spectrum matches (PSMs) for viral proteins by the sum of all PSMs in the sample in B05 (a and b) and B01 (c and d). a) and c) represent samples from the cellular metaproteome, b) and d) represent samples from the extracellular metaproteome. vMAGs that are among the top 30 most abundant across all samples are colored. The dashed green lines represent the average cell count for each time point.

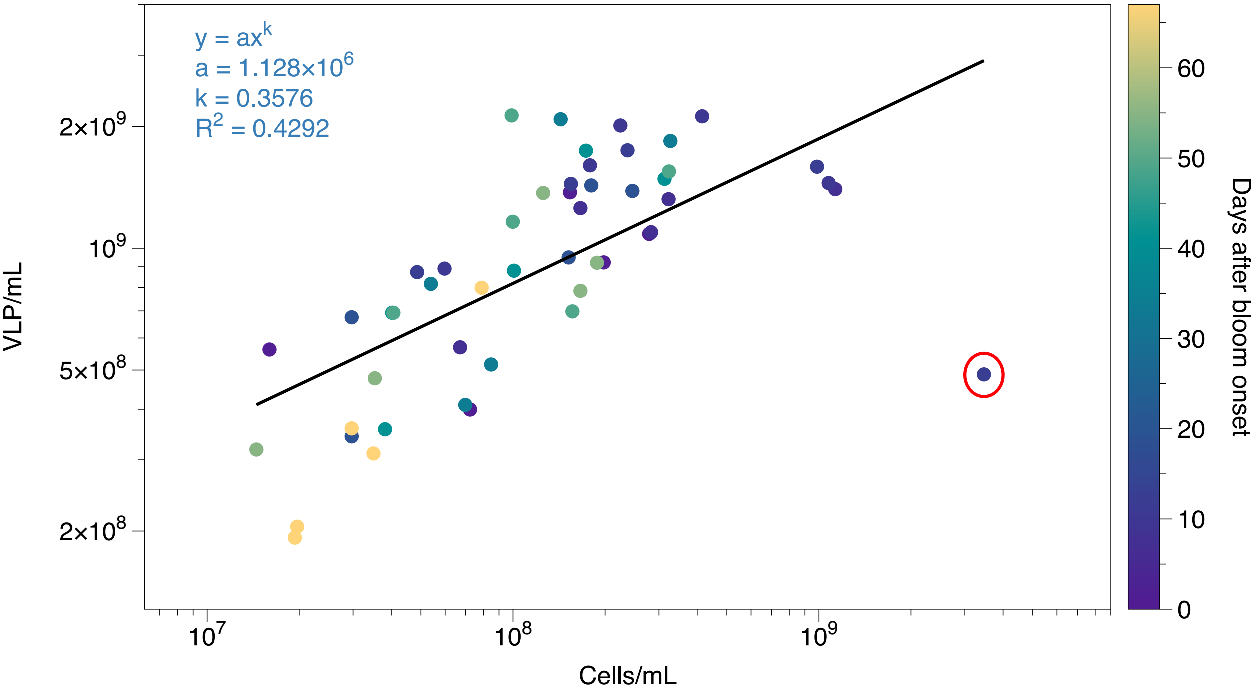


**S9. Cell counts from 2024 vs VLP counts from 2024 fit by a power law.** Points are colored by the number of days after bloom onset. The point with the highest cellular abundance is circled in red. With outliers (>2 standard deviations from the mean, n = 2) removed from the calculated fit, R2 increases to 0.6549 (not shown).

#### Supplementary methods

##### Sample collection

Samples were collected with a custom sampling pole as described in [1]. For each depth sampled, we first flushed the tubing inside the sampling apparatus by drawing and discarding 50 mL using a syringe. We then connected an acid washed, flushed (3x with Argon), and evacuated anaerobic media bottle to the sampling port. The liquid from the water column filled the evacuated media bottle because of the pressure differential. Samples were stored and transported chilled until processing, and were processed within 3 hours of collection.

##### Direct counts

For cell counts, we fixed 1 mL of sample in a final concentration of 2% formaldehyde in 1× PBS for 1 hour. In 2021, we stored cell fixed samples at 4 °C until counting. In 2024, we stored samples at -80 °C until counting.

To prepare samples for cell counts, we followed the protocol from [1]. We diluted samples to a final concentration of ~$4 \times 10^5$ cells $mL^{-1}$ in 1× PBS, then sonicated them to disrupt aggregations of cells with a probe sonicator (Sonics Vibra-Cell™, VC 505) at 21% amplitude with 20s on, 20s off pulses of sonication for a total duration of 4 minutes. We filtered the sonicated samples using a glass filter tower onto a 0.2 µm polycarbonate filter (MilliporeSigma, GTTP04700). We added 1 ml of 1× PBS to the filter tower prior to the addition of the sample to prevent the meniscus from concentrating particles towards the edges of the filter and rinsed the filter tower with an additional milliliter of 1× PBS after sample filtration. The filter tower was briefly acid washed in 1 M HCl and rinsed with DI water between samples. Once the filters were dry, we stained them with DAPI (4',6-diamidino-2-phenylindole, 1 µg $ml^{-1}$) for 10 minutes, dried them again, and mounted them onto slides with an anti-fade mounting solution of Citifluor (Fisher, 50-302-34) and Vectashield (Fisher, NC9265087) at 4 to 1 ratio.

For VLP counts, we pre-filtered 1 mL of sample using a 0.2 µm syringe filter (Whatman™ Uniflo™, sterile PES, WHA9915-2502), then fixed the filtrate in a final concentration of 2% formaldehyde in 1× PBS. Samples were stored at -80 °C until counting.

To prepare samples for VLP counts we diluted and sonicated samples as described for cell counts. Following [2], we filtered sonicated samples using a glass filter tower onto a 0.02 µm Anodisc filter (Whatman, WHA68096002) with a 0.45 µm support filter (Sartorius, 14555400) with 1 mL 1× PBS added before and after the sample as above. We stained dried filters with Sybr Green I (Thermo Fisher, S7563) at a 1:400 dilution of the 10000× stock for 25 minutes. We mounted filters onto slides with an anti-fade mounting solution of *p*-phenylenediamine dihydrochloride (Sigma, P-1519) in 1:1 PBS:Glycerol (0.1% f.c.).

Filters were visualized on a Zeiss Axiolab epifluorescence microscope with appropriate fluorescent filters. For each filter, we counted twenty 10 by 10 grids and calculated counts per mL as follows:

$$\frac{Filterable\ area}{Area\ counted} \times number\ of\ particles\ counted \times \frac{1}{volume\ filtered\ (mL)}$$

Where the filterable radius is the radius of the filter tower, and the area counted represents the area of the grid at a magnification of 1000×. The volume filtered represents the total amount of sample filtered with dilutions taken into account.

**Catalyzed reporter deposition fluorescence in situ hybridization (CARD-FISH)**

CARD-FISH was performed as in [1] (see https://www.protocols.io/view/catalyzed-reporter-deposition-fluorescence-in-situ-rm7vzjebrlx1/v1 for a detailed protocol). We fixed and filtered samples onto 0.2 µm polycarbonate filters (MilliporeSigma, GTTP04700) with a 0.45 µm nitrocellulose support filters (Sartorius, 14555400) as above, except that fixed cells were washed once with 1× PBS, then stored in 1:1 ethanol:1× PBS at -20 °C. We used two probes: GSB-532, targeting green sulfur bacteria [3] and NON338, a nonsense probe [4]. We used a lysozyme solution of 1000 kU $ml^{-1}$ in 0.05 M EDTA [pH 8.0], 0.1 M Tris-HCl [pH 7.5], incubated for 1h at 37 °C to permeabilize cells, then washed filters in ultrapure water. To inactivate endogenous peroxidases, we incubated filters with 0.01 M HCl for 15 minutes at room temperature, followed by washing with ultrapure water and 96% ethanol (EtOH). We air dried filters prior to hybridization with HRP-labeled probes. We used a 10% w/v sterile blocking solution with blocking reagent (Roche, 11096176001) in maleic acid buffer (100 mM maleic acid and 150 mM NaCl [pH=7.5]). We combined 300 μl of hybridization buffer (0.9 M NaCl, 20 mM Tris-HCl [pH 7.5], 10% w/v dextran sulfate, 0.02% w/v sodium dodecyl sulfate (SDS), 1% w/v blocking reagent, and 10% formamide (GSB-532) or 35% formamide (NON-388) with 1 μl of HRP-probe working solution (50 ng $\mu l^{-1}$; 0.16 ng $\mu l^{-1}$ final), and incubated filters at 46°C for 2-3 h. We washed filters with buffer (0.45 M NaCl (GSB-532) or 0.08 M NaCl (NON-388), 5mM EDTA [pH 8.0], 20mM Tris-HCl [pH 7.5], 0.01% w/v SDS) for 10 minutes at 48°C and in 1× PBS for 15 minutes at room temperature. We transferred filters to amplification buffer (2 M NaCl, 0.1% w/v blocking reagent, 0.0015% H2O2, 10% w/v dextran sulfate; in 1× PBS) containing Alexa Flour 488 (Invitrogen, B40957) tyramide (1 mg $ml^{-1}$) in a ratio of 1:500 and incubated them in the dark for 30 minutes at 46 °C. We incubated filters in 1× PBS for 10 minutes at room temperature in the dark and washed them in ultrapure water and 96% EtOH. To visualize cells, we mounted filters in an antifade mounting medium (Citifluor:Vectashield, 4:1) on a microscopy slide.

**DNA extraction**

Sample preparation and DNA extraction was carried out as described in [1]. A detailed protocol for DNA extraction is available at https://www.protocols.io/view/dna-extraction-with-ctab-and-chloroform-isoamyl-al-14egn7b9yv5d/v1.

We added 1 mL of DNA extraction buffer (DEB; 100 mM Tris-HCl (pH 8), 100 mM EDTA (pH 8), 100 mM sodium phosphate buffer (pH 8), 1.5 M NaCl, 1% CTAB) to an Eppendorf tube containing a filter. To mechanically disrupt cells, we performed three freeze-thaw cycles (-20 °C to 55 °C). For chemical lysis, we added lysozyme (2 mg/mL f.c.) and incubated the sample at 37 °C for 30 min, then added Proteinase K (0.2 mg/mL f.c.) and incubated the sample at 37 °C for 30 min. Following this incubation, we added sodium dodecyl sulfate (SDS) (1% f.c.) and incubated samples at 65 °C for 2 h with inversion every 30 minutes. To phase separate the DNA fraction of the lysate, we twice mixed the sample with chloroform-isoamyl alcohol (24:1, pH 8), and centrifuged it to separate the DNA-containing aqueous fraction, which we collected. To precipitate DNA, we added 100% isopropanol at 0.6 × sample volume and incubated the sample for 2 hours at room temperature. We centrifuged the sample 20,000 × rcf for 30 minutes to collect the precipitated DNA in a pellet. We washed this pellet twice with ice cold 70% ethanol, briefly air dried it, and resuspended it in 100-250 µL of nuclease-free water. To assess DNA quality and quantity before sequencing, we used a Qubit 2.0 fluorometer and NanoDrop 2000 spectrophotometer.

**Oxford Nanopore Sequencing**

We sequenced a single sample of extracted DNA (>0.2 µm size fraction) from September 1st, 2021 with Oxford Nanopore Technologies (ONT) ligation sequencing kit (SQK-LSK110) following the manufacturer’s instructions. We prepared a total of 1.06 µg of high-molecular weight gDNA as per manufacturer's protocols. We conducted DNA repair and end-preparation to generate blunt-ended, dA-tailed fragments. We used the ligation module to ligate sequencing adapters, and purified adapter-ligated DNA using AMPure XP beads, including a long fragment buffer (LFB) wash to enrich for high–molecular weight fragments, and incubated at 37 °C to improve recovery. We eluted the final library in elution buffer and quantified using Qubit HS (Thermo Fisher, Q33231) prior to sequencing.

We loaded the resulting library (8.9 ng/µL) onto the flow cell (FLOW-MIN106; FA56891) and sequenced with a MinION sequencer (Mk1B; Oxford Nanopore) following the SQKLSK110 manufacturer's protocol. We ran MinKNOW (v 5.1.8) following the Mk1B sequencing protocol with adaptive mode on and live-base calling turned, resulting in raw FAST5 format files for post-run basecalling.

**Nanopore Sequence Data Processing**

We processed Nanopore sequence data following [5] with recommendations for assembling bacterial genomes from Oxford Nanopore sequencing with short-read Illumina polishing. To convert raw fast5 format files to pod5 format we used the pod5 (v0.3.6) python module (Oxford Nanopore Technologies, ONT). We used Dorado (v0.5.3, ONT) for basecalling, the quality filtered the reads using Fitlong (v0.2.1) [6] without an external reference with the following settings: min_length=1000, keep_percent=90, target_bases=500000000. For long-read assembly we used Flye (v2.9.3) using the optional argument for corrected ONT reads (nano-corr) [7]. We used Medaka (v1.6.0, ONT) for sequence correction. To polish long-read sequence data with short reads from the same sample from our Illumina dataset (see below), we used Polypolish (v0.6.0) [8].

Given recent concerns about the quality of long read assemblies [9], we checked the quality of the genome using the following: 1) manual read mapping inspection to assess evenness of coverage and any regions of low to zero coverage, 2) manual inspection of GC coverage graphs to detect sharp changes, 3) a length comparison with other closely related genomes, 4) assessment of long read clipping events using *anvi-script-find-misassemblies* [10]. Using these approaches, we failed to detect anomalous regions suggestive of misassembly.

**Illumina sequencing**

Shotgun metagenomics was performed by Psomagen, Inc, as described by [1]. DNA quality screening was performed using Picogreen (Thermo Fisher) and the genomic DNA ScreenTape assay (Agilent). Illumina's DNA Prep Kit was used to prepare libraries by mixing samples with bead-linked transposomes and incubating at 55°C for 15 minutes to fragment and tag DNA with adapter sequences. PCR was performed on tagmented samples using a limited-cycle PCR program with indexes from Illumina's Nextera DNA Unique Dual Indexes kit. PCR products were purified with magnetic beads and quality control was performed using the D5000 ScreenTape assay (Agilent) and Picogreen (Thermo Fisher). Highly concentrated libraries were normalized to 5 nM. Approximately 1.5-2 nM of libraries were loaded onto the flow cell and sequenced using a NovaSeq6000 S4 (v1.5) platform with PE150 chemistry (paired-end sequencing, 150 bp).

**Short read metagenomic data processing and metagenome assembled genome (MAG) generation**

We performed read quality control for MAG generation and read-mapping as described in [1]. We removed adapters using trimmomatic (v0.36) for paired-end reads (ILLUMINACLIP:2:30:10, LEADING:3, TRAILING:3, SLIDINGWINDOW:4:15, MINLEN:65) [11]. To remove phiX control reads, we mapped reads against phiX174 (NCBI NC_001422) and removed the reads that matched. We removed low quality reads using iu-filter-quality-minoche from illumina-utils (v2.12) [12]. To validate quality control, we assessed read quality with FastQC (v0.11.4) [13].

We performed assemblies separately for each time point (with filters from each size fraction combined) and co-assemblies combining all time points in each bloom using MEGAHIT (v1.2.9) [14] with contigs shorter than 1000 bp removed with anvi-script-reformat-fasta [10]. To generate MAGs, we combined the bins identified by four different binning tools: MaxBin2 (v2.2.7) [15], MetaBAT2 (v2.12.1) [16], CONCOCT (v1.0.0), and SemiBin (v1.4.0) [17], which uses a deep siamese neural network. We used metaWRAP to index assemblies and map reads with BWA [18] and perform binning with the first three binning tools. To find the optimized, non-redundant set of bins from the combined binning output from all four binning algorithms, we used DASTool (v1.1.6) [19].

**Identification of viral sequences and viral metagenome assembled genome (vMAG) generation**

We performed read quality control for viral assemblies by first removing low quality reads using iu-filter-quality-minoche from illumina-utils (v2.12) [12], then removing adapters using trimmomatic (v0.36) for paired-end reads with the following settings ILLUMINACLIP:adapters.fa:2:30:10 LEADING:3 TRAILING:3 SLIDINGWINDOW:4:15 MINLEN:75. We checked for the presence of phiX control reads in quality controlled read files using Bowtie[20]2 v2.5.1. No reads maps to the phiX reference. To decrease the likelihood of chimeric viral assemblies and increase the yield of viral contigs, we combined quality-controlled reads by size fraction across all time points for the two smallest size fractions, 0.025-0.1 µm and 0.1-0.22 µm from 2021. Reads from 2024 and 2025 were not included in the co-assemblies. We co-assembled each pool with MEGAHIT v1.2.9 [14] ( --presets meta-sensitive --min-contig-len

1000). We identified viral contigs in the co-assemblies with geNomad v1.8.1 [21] and VirSorter2 v2.2.4 [22] (--min-length 1000 --min-score 0.5). We combined the output of these two tools and dereplicated it with seqkit rmdup v0.16.0 [23]. To assess the quality of these putatively viral contigs, we used CheckV v1.0.1 [24] and manually filtered the contigs using standards modified from [25]. We removed a) all contigs where viral gene = 0 and host gene > 0 as identified by CheckV, and b) contigs shorter than 5 kbp. Following the approach of [26], we dereplicated these quality controlled viral contigs using CD-HIT v4.6 [27] with the following parameters *cd-hit-est -M 20000 -c 0.95 -aS 0.85*. We binned dereplicated viral contigs with vRhyme v1.1.0 [28], excluding contigs identified by CheckV [24] as being complete or prophages. We linked contigs in bins using link_bin_sequences.py script (provided by vRhyme) for downstream analyses.

To create the initial vMAG database, we combined binned vMAGs with contigs representing complete viruses (as identified by CheckV), prophages, and unbinned viral contigs. We assessed this vMAG database with CheckV [24] again and only vMAGs longer than 10 kbp and classified as "High quality" or "Complete" were used in downstream analyses. We used vRhyme [28] to dereplicate this final vMAG database with the following settings: –derep_only –method longest, which only considers length in dereplication. The final vMAG database contained 2750 dereplicated high quality vMAGs.

We assigned viral taxonomy of the final vMAG database with geNomad [29]. To further classify the vMAGs classified as *Megaviricetes* (a class of giant viruses), we filtered the MAGs to a minimum contig length of 100 kb and ran Viralrecall v3.0 (https://github.com/abdealijivaji/ViralRecall_3.0) [30]. We manually inspected the output of Viralrecall based on contig length ( >100 kbp) and giant virus markers. We then classified selected vMAGs at the Order and Family level with TIGTOG [31]. We manually inspected all other viruses classified as non-*Caudoviricetes* to confirm their classification. Any vMAG with a geNomad classification that was not supported upon visual inspection was instead classified as "Unknown". We used Cenote-Taker 3 v3.0.0 [32] for functional annotations.

To determine relative abundances of both MAGs and vMAGs we used CoverM v0.6.1 [33]. Reads from all three years, 2021, 2024, and 2025 were mapped to MAGs and vMAGs separately.

**vMAG host identification**

We used iPHoP v1.3.3 [34] with the Aug_2023_pub_rw database modified to include the prokaryotic bin database described above (yet un-dereplicated) to identify putative hosts of vMAGs. iPHoP integrates host-based approaches (sequence homology, CRISPR spacer matches, and k-mer frequency matches) with phage-based approaches (the comparison of an input phage sequence with a database of phages with known hosts) using a random forest classifier to obtain an integrated confidence score (iPHoP-RF) for each phage-host match.

**Metaproteomics**

***Cell lysis, sample cleanup, and sample preparation***

To capture the cellular and extracellular protein fractions, we separately extracted protein from cells collected on 0.22 μm polyethersulfone Sterivex filter (cellular) and supernatant from 100 mL of sample centrifuged at 20,000 × rcf for 15 minutes at 4 °C (extracellular). Each of these sample types required a different cell lysis and extraction method.

Sterivex filter units were thawed and the filters removed from the housings. We extracted biomass from Sterivex filters and lysed cells using a method adapted from [35]. We divided the filter into approximately 1 $cm^2$ pieces with a sterile scalpel and suspended the pieces in 1.7 ml of lysis buffer (4% (w/v) SDS, 100 mM Tris-HCl pH 8.0). We heated samples to 97°C for 15 minutes, incubated them for 1 hour at room temperature, and then briefly vortexed them for 1 minute. We centrifuged the samples at 10,000 × g for 5 minutes and transferred the supernatant to a new tube. We added 1.7 ml of fresh lysis buffer to the filter pieces, vortexed and centrifuged as before, and combined the resulting supernatant with the previous supernatant for the final lysate.

For the extracellular samples we concentrated supernatants of pelleted bloom samples (~12 ml) on Amicon Ultra-15 (10 kDa cutoff; Millipore Sigma) filtration units by centrifuging at 4,000 × g for 30 minutes at 4 °C to a volume of 300-400 µl. We added lysis buffer at a 1:2 ratio (sample:lysis buffer) and heated samples to 95 °C for 10 minutes for the final lysate.

To remove interfering environmental compounds (e.g., humic acids) from all samples, we subjected samples to an overnight trichloroacetic acid (TCA) precipitation based on a previously described method [36]. We suspended the samples in 10% TCA and stored them at -80 °C overnight. We thawed the samples and centrifuged at 21,000 × g for 15 minutes at 4 °C to pellet the precipitated protein. The resulting pellet was washed in 1 ml of ice cold 100% acetone, vortexed briefly and centrifuged at 21,000 × g for 15 minutes at 4 °C. The final pellets were air dried and resuspended in 60-120 µl of SDT-lysis buffer (4% (w/v) SDS, 100 mM Tris-HCl pH 7.6, 0.1 M dithiothreitol (DTT)).

We prepared samples for LC-MS/MS analysis using a modified filter-aided sample preparation method [37]. The samples in SDT-lysis buffer were heated at 95 °C for 10 minutes before being loaded onto 10 kDa PES membrane centrifugal filters (VWR International) with UA buffer (8 M urea, 0.1 M Tris-HCl pH 8.5) at a ratio of 60 µl of sample lysate to 400 µl of UA buffer. We centrifuged filter units at 14,000 × g for 20 minutes. We washed the filter units by adding 200 µl UA buffer and centrifuging as before. We added 100 µl of IAA solution (0.05 M iodoacetamide in UA solution) to the filter, mixed at 600 rpm for 1 minute (Benchmark Scientific Inc., model: H5000-HC), and incubated at 22°C for 20 minutes. We washed the filters three times with 100 µl UA before performing a buffer exchange to 50 mM ammonium bicarbonate (ABC buffer) with three washes of 100 µl ABC buffer. We digested proteins overnight by adding 0.8 to 1 µg of MS grade trypsin (Thermo Scientific Pierce) in 40 µl of ABC buffer, mixing at 600 rpm for 1 minute, and incubating overnight at 37 °C in a wet chamber. After, we eluted peptides by centrifuging the filter units at 14,000 × g for 20 minutes, followed by an additional elution with 50 µl of 0.5 M NaCl, mixed at 600 rpm for 1 minute and centrifugation as before. We estimated the peptide concentrations of final eluates using the Pierce Micro BCA assay (Thermo Scientific Pierce) following the manufacturer instructions.

***1D nanoflow liquid chromatography and tandem mass spectrometry (LC-MS/MS)***

For each sample 2000 ng of peptide were separated using an UltiMate 3000 RSLCnano UHPLC system (Thermo Fisher Scientific). Samples were first loaded onto a 5 mm, 500 µm C18 Acclaim PepMap100 pre-column (Thermo Fisher Scientific) for desalting, prior to separation on a 75 µm x 75 cm EASY-spray column packed with PepMap RSLC C18, 2 µm material (Thermo Fisher Scientific) at 60°C. We separated peptides using a previously published 140 minute reverse-phase gradient [38]. The eluting peptides were ionized via electrospray ionization (ESI) with an Easy-Spray source and measured using an Orbitrap Exploris 480 Mass Spectrometer (Thermo Fisher Scientific) by a data dependent acquisition [38]. In brief, the precursor scans were acquired for a 380 to 1,600 m/z window at a resolution of 60,000 with a maximum injection time of 200 ms and a normalized AGC target of 3e6. The 15 most abundant ions were isolated for $MS^2$ analysis (Top15), using a dynamic exclusion of 25 s. $MS^2$ spectra were acquired at a maximum injection time of 50 ms and an AGC target of 1e5. Isolated ions were subjected to a normalized collision energy of 27% in the HCD cell prior to measurement in the Orbitrap mass analyzer at a resolution of 15,000. An average of 81,000 spectra collected per sample.

***Database construction and protein identification***

We created a protein sequence database using predicted protein sequences from the MAGs, vMAGs, and protein sequences predicted from unbinned contigs. We constructed this database following previous recommendations [39], reducing redundancy using CD-HIT for sequence clustering, and prioritizing taxonomically resolved sequences from MAGs over sequences from unbinned contigs with CD-HIT-2D [27, 40]. To account for the presence of common laboratory contaminants, we added the cRAP database (https://www.thegpm.org/crap/). The final 1,253,648 sequence database is available in the PRIDE repository along with the raw spectral data.

We identified peptides and inferred proteins from this data by searching the $MS^2$ spectra against the custom database using Proteome Discoverer version 2.3 (Thermo Fisher Scientific) using a previously described method [41]. Proteins were quantified using peptide spectral counts (PSMs) and we only retained proteins identified with a false discovery rate (FDR) of less than 5% that were designated as master proteins. We used the spectral count data to calculate normalized spectral abundance factor percentages (NSAF%) and re-normalized the NSAF% values of the primary organism in the bloom, *Prosthecochloris sp.* and its associated plasmid (contig_20) to obtain organism normalized NSAF% values (orgNSAF%)  [42–44].

To assess the percent biomass contribution from the viral community, we used a modified protocol from [45]. We filtered the metaproteomic database for proteins with 5% FDR confidence and at least one protein unique peptide (note the more lenient standards as compared to [45]), then divided the PSM of each protein by the total PSM for each sample to calculate percent biomass. Percent biomass contributions for each protein were summed by organism to compare biomass contributions.

#### Broader analysis of bloom forming organisms

The full list of aquatic environments from Sandpiper included in this analysis is as follows: "lake water metagenome", "brine metagenome", "aquatic metagenome", "riverine metagenome", "groundwater metagenome", "salt lake metagenome", "pond metagenome", "estuary metagenome", "soda lake metagenome", "marsh metagenome", "marine metagenome", "freshwater metagenome", "aquifer metagenome", "mangrove metagenome", "hypersaline lake metagenome", "salt marsh metagenome", "seawater metagenome", "wetland metagenome", "marine plankton metagenome".

### Supplementary results

#### Viral metagenome assembled genomes (vMAGs) are not bioinformatically linked to GSB-TRL01

A total of 2750 viral metagenome assembled genomes (vMAGs) were identified. Of these, the majority were *Caudoviricetes* (supplementary figure S2). Of the non-*Caudoviricetes*, 37 vMAGs were identified as *Megaviricetes*, two as *Maveriviricetes* (virophages), one as *Faserviricetes* (ssDNA viruses), seven as *Bamfordvirae* without a class-level identification, and 32 as unknown viruses.

Most vMAGs were not successfully matched to a host (figure 3b, S2) with iPHoP. Although iPHoP integrates multiple host and phage based approaches, and outperforms other sequence-based tools [34], host-phage matching based on sequences alone has several drawbacks. Sequence homology between phages and hosts does not always occur between phage-host pairs. The integration of CRISPR spacers may not yet have occurred, or phages may have evolved to escape targeting by integrated spacers. Approaches based on the "guilt by association" of phage sequences to the sequences of phages with known hosts are severely limited by the lack of representation of phages infecting certain taxa in databases (the Chlorobiota, for example, have no isolated phage representatives).

Of those phages matched to hosts, the majority were *Caudoviricetes.* The most common host phyla were: *Pseudomonadota* (80), *Bacteroidota* (43) and *Thermoplasmatota* (20) (figure 3c, S2). None were matched within the *Chlorobiota* (*Bacteroidota_A* by GTDB taxonomy). Distinct relative abundance shifts occurred over the time series (figure 3c). Although none of the vMAGs were linked to *Chlorobiales*, the influence of the bloom on the viral community is demonstrated by the clear relative abundance shift between samples during the bloom in 2021 (2-8 days after the start of sampling), and the sample from after the bloom (58 days after the start of sampling; figure 3c). For example, phages linked to the *Pseudomonadales* and the *Izemoplasmatales* became less relatively abundant, whereas phages linked to the *Enterobacterales*, *Enterobacterales_A*, and *Flavobacterales* became more relatively abundant.

In the 2024 and 2025 samples, the relative abundance of the host matched vMAGs is very low. This likely reflects the rapid turnover of the viral community, since the vMAGs were generated using data from 2021 only. The number of reads mapped to vMAGs in general (irrespective of host ID) was not diminished as compared to the larger size fractions from 2021 in these samples (figure 3b).

The 0.02-0.1 μm size fraction has a larger proportion of viral reads than the larger size fractions, indicating that the size fractionation treatment successfully selected for viral DNA (figure 3bc).

**GSB-TRL01's genome contains a putative phage satellite**

The putative phage satellite is located in a transposon-rich region of the genome (figure 4a) and has lower coverage than the surrounding region of the genome (figure S4), indicating that it may only be present in a subset of the population. The putative satellite is ~14kb long and was classified by SatelliteFinder [46] as a phage inducible chromosomal island (PICI) Type C variant 1 satellite. Type C PICIs have three of five core PICI components—variant 1 PICIs have an integrase, a regulatory gene (usually alpA), and a DNA primase/replicase. Our phage satellite additionally contained a DNA primase, a complete Type I restriction-modification system, and a possible fragment of a CBASS system. It is unknown whether Type C PICIs are functional satellites or represent defective PICI remnants, or even degraded prophages [46].

**Supplementary discussion**

**Integrase detection in the proteome does not necessarily reflect active lysogeny**

Besides the explanations outlined in the main text, another possible explanation for the reduction in free VLPs at the height of the bloom is the "Piggyback-the-Winner" hypothesis, which posits that lysogeny becomes prevalent at high cell densities, possibly because of superinfection dynamics [47]. We do not see evidence for lysogen integration, as no lysogens were detected in our genome from the height of the bloom. We did detect an integrase with relatively high expression in the metaproteome, but this integrase (belonging to vRhyme_bin_17) has 95% amino acid percent identity with the integrase belonging to the phage satellite located on the GSB-TRL01 genome (figure 4a). Other proteins on this vMAG do not have high expression in the proteome, so we believe the detection of this integrase reflects expression from the phage satellite, rather than this vMAG. The high percent identity between these two integrases may reflect recent genetic exchange between the phage satellite and the vMAG—possibly identifying this vMAG as the helper phage of the phage satellite.

**There is evidence for active phage infection during the bloom**

Several vMAGs had higher read coverage during the bloom (supplementary figure S3). A major capsid protein from vRhyme_bin_7, a viral population genome unmatched by iPHoP to a host, is present in the metaproteomes of both the cellular and extracellular proteomic fractions during the bloom. This pattern suggests active lytic infection, since proteins from actively replicating viruses are expected to appear in the cellular fraction, while free viral particles are expected to appear in the extracellular fraction. vRhyme_bin_7 is an especially strong candidate for a GSB-TRL01 infecting phage since its abundance in both the metagenome and in the proteome follow cellular abundances of GSB-TRL01. The increase in abundance of proteins, mostly structural, from several other vMAGs in the metaproteome of the extracellular fraction during the bloom (supplementary figure S7) further suggests that infection does occur, albeit not on a large scale.

One possibility is that these phages are not GSB-TRL01 specialists and are instead opportunistic generalist phages. This aligns well with the idea of "enhanced infection" [48], wherein resistance against certain phages opens bacteria up to infection by different phages. This may represent an effective infection strategy for the phage, given that GSB-TRL01 is rare for most of the year. Supporting this hypothesis, CRISPR spacer analysis suggests that many cyanophages infecting bloom forming cyanobacteria have broad host ranges [49]. This indicates that bloom forming organisms may more commonly be infected by generalist phages, an idea that merits future investigation.

**The abundance of defense mechanisms in GSB-TRL01's genome are likely to exert costs of resistance**

CRISPR-Cas systems seem to be associated with both a constitutive and an inducible cost [50]. There is therefore a diversity of regulatory mechanisms that control the expression of CRISPR-Cas systems. While the conditions of up and down regulation of these systems appear to be niche specific, CRISPR-Cas regulation in response to environmental or cellular conditions is widespread. For example, Cas genes related to spacer acquisition have been shown to be upregulated via quorum sensing, leading to increased spacer acquisition in higher density microbial communities [51]. A proteomic study in *Streptococcus thermophilus* infected by phage 2972 found that the expression of some Cas proteins dramatically increased over the course of infection and increased with increasing multiplicity of infection, indicating that the expression of Cas proteins dynamically responds to infection pressure [52]. The expression of Cas genes from the CRISPR-Cas systems both of the GSB-TRL01 genome and of the plasmid at levels as high as 0.1% (orgNSAF%) therefore suggests the active use of these systems.

Restriction modification systems are costly both because of autoimmunity risks and because of energy expenditure. The GSB-TRL01 genome encodes four Type I restriction modification (RM) systems (one of which is contained within the putative phage satellite; figure S4) and the plasmid encodes an additional two Type II RM systems. RM systems can also be extremely energetically costly (the Type I RM enzyme *Eco*R124I consumes one ATP per modified base-pair [53]), likely leading to the regulation of their expression [50]. Indeed, a transcriptional regulator is found between two methyltransferase genes in the genome's defense island. RM systems also represent a cost to the host in the form of the risk of autoimmunity, especially under nutrient limited conditions [54].

Abortive infection (Abi) systems have been shown to incur a fitness cost, but the mechanism is unknown and may result from either expression costs of the system itself, or from erroneous triggering of cell death [55]. Since the overexpression of the toxin can lead to cell death, Abi systems are tightly regulated [56]. There are many possible mechanisms for regulation in these systems, but the detection of toxins from GSB-TRL01's Abi systems in the metaproteome (figure 5) may indicate active abortive infection.

The other defense systems encoded on GSB-TRL01's genome are likely also costly, but the mechanisms of these costs and the regulation of the systems have not been described. The expression of the anti-plasmid Wadjet system (figure 5), for example, may be regulated by homologs of the protein brxR, which regulates the BREX system [57]. This regulation implies that it is only advantageous to express the system in some situations and that its expression may indicate active use.

**The maintenance of a large conjugative plasmid contributes to anti-phage defense**

The acquisition of plasmids is costly for bacterial hosts, and although the host can eventually adapt to compensate for some costs, the expression of genes found on the plasmid (especially for large complexes like Type IV secretion systems) will continue to represent an energetic burden [58]. T4SS have diverse regulation systems: expression is either induced in response to environmental factors, or tightly regulated to avoid exerting too heavy of a cost on the host cell [59]. While the regulation mechanisms employed by this plasmid are unknown, the T4SS contained what appears to be a repressor protein, which may indicate that it represses the expression of its conjugation machinery when unused. The presence of conjugative machinery—including proteins forming the core conjugative complex and involved in pilus formation—therefore suggests that plasmid transfer by conjugation was actively occurring in the GSB-

TRL01 population. This relatively large conjugative plasmid that is maintained in a majority of the population and appeared to be undergoing active transfer likely provides substantial fitness benefits. Since 50% of the annotated genes on the plasmid are related either to conjugation or to anti-mobile element defense, these fitness benefits may derive from the plasmid's anti-mobile genetic element defense systems.

Systems used for anti-mobile element defense, such as toxin anti-toxin systems and restriction modification systems, can also double as "addiction mechanisms" which insure plasmid maintenance by killing daughter cells that fail to inherit it [60]. Competition between mobile elements is a major driver of accumulation of defense systems [61]. Some types of anti-phage defense systems, such as CRISPR-Cas, are often found on mobile genetic elements like plasmids. CRISPR-Cas systems encoded on MGEs may work synergistically or antagonistically with host defense systems [62]. Other types, such as RM systems, are relatively uncommon on plasmids, although RM systems are more common plasmids with conjugative machinery [63].

Interestingly, GSB-TRL01 encodes a complete Wadjet anti-plasmid system, which cleaves closed circular DNA to prevent plasmid transformation [64]. The presence of a conjugative plasmid within a cell encoding the Wadjet system is surprising given that Wadjet defends against plasmids, but the plasmid's large size (92 kbp) may suppress cleavage because the upper size limit for topology detection by Wadjet seems to be between 50-100 kbp [65].

**Investment in defense as an ecological trait of bloom forming organisms**

As outlined in the main text, bloom-forming organisms are generally thought to be opportunistic organisms that grow to dominate under advantageous nutrient or environmental conditions. In theory, they align with the ecological terms (used variously in different subfields): zymogenous organisms (from soil ecology), copiotrophs (from marine ecology), and r-strategists (characterized by the production of large numbers of offspring) [66]. They are likely to have a fast growth rate under optimal conditions, have a high nutrient demand and low substrate use efficiency, and variable population dynamics. Importantly, not all organisms classified as zymogenous, copiotrophic, or r-strategists will form the dense, low diversity blooms that are investigated here. A recent study sought to understand the traits of bloom forming organisms that sets them apart from other copiotrophs and found that they have a high codon usage bias and are enriched in signal transduction mechanisms and transcription related genes [67]. Few other studies have sought to experimentally generalize the qualities that make bloom formers successful. We show that many have the genomic capabilities to be defense specialists, as outlined by Thingstad in the Kill-the-Winner hypothesis [68], and suggest that this may be a generalizable trait of bloom formers.

12. merenlab/illumina-utils. 2025. Meren Lab, 2025.

13. Babraham Bioinformatics - FastQC A Quality Control tool for High Throughput Sequence Data. https://www.bioinformatics.babraham.ac.uk/projects/fastqc/. .

43. Kleiner M. Normalization of metatranscriptomic and metaproteomic data for differential gene expression analyses: The importance of accounting for organism abundance. PeerJ Inc., 2017.
